# Skewed distribution of spines is independent of presynaptic transmitter release and synaptic plasticity and emerges early during adult neurogenesis

**DOI:** 10.1101/2023.03.15.532740

**Authors:** Nina Rößler, Tassilo Jungenitz, Albrecht Sigler, Alexander Bird, Martin Mittag, Jeong Seop Rhee, Thomas Deller, Hermann Cuntz, Nils Brose, Stephan W. Schwarzacher, Peter Jedlicka

**Author notes:** Corresponding Authors: Nina Rößler ICAR3R - Interdisciplinary Centre for 3Rs in Animal Research Faculty of Medicine Justus-Liebig-University Rudolf-Buchheim-Str. 6 D-35392 Giessen Peter Jedlicka ICAR3R - Interdisciplinary Centre for 3Rs in Animal Research Faculty of Medicine Justus-Liebig-University Rudolf-Buchheim-Str. 6 D-35392 Giessen. joint first authors (Nina Rößler, Tassilo Jungenitz).

## Abstract

Dendritic spines are crucial for excitatory synaptic transmission as the size of a spine head correlates with the strength of its synapse. The distribution of spine head sizes follows a lognormal-like distribution with more small spines than large ones. We analysed the impact of synaptic activity and plasticity on the spine size distribution in adult-born hippocampal granule cells from rats with induced homo- and heterosynaptic long-term plasticity in vivo and CA1 pyramidal cells from Munc-13-1-Munc13-2 knockout mice with completely blocked synaptic transmission. Neither induction of extrinsic synaptic plasticity nor the blockage of presynaptic activity degrades the lognormal-like distribution but changes its mean, variance and skewness. The skewed distribution develops early in the life of the neuron. Our findings and their computational modelling support the idea that intrinsic synaptic plasticity is sufficient for the generation, while a combination of intrinsic and extrinsic synaptic plasticity maintains lognormal like distribution of spines.

## Introduction

A variety of features in the brain including dendritic spine size (Loewenstein et al., 2011; Montero-Crespo et al., 2020; Santuy et al., 2018), synaptic strength (Cossell et al., 2015; Ikegaya et al., 2013; Lefort et al., 2009; Song et al., 2005) and neuronal firing rate (Mizuseki & Buzsáki, 2013) are strongly positively skewed with a heavy tail, displaying a lognormal-like distribution. Lognormal-like distributions of synaptic and firing rate parameters are thought to play a fundamental role in the structural and functional organization of the brain (Barbour et al., 2007; Buzsáki & Mizuseki, 2014; Kasai et al., 2021), and a number of explanations for the emergence of such distributions in active and plastic networks have been proposed.

Spines are plastic and motile structures of neuronal dendrites that function as postsynaptic sites for excitatory inputs. The spine head contains the postsynaptic density (PSD) with AMPA and NMDA glutamate receptors (Ziff, 1997). The size of the PSD correlates with spine head size, the number of presynaptic vesicles (Harris et al., 1992; Harris & Stevens, 1989), and the density of postsynaptic receptors (Matsuzaki et al., 2001; Nusser et al., 1998; Takumi et al., 1999; Zito et al., 2009). Therefore, spine head size has been used as a morphological proxy for synaptic strength (Asrican et al., 2007; Bromer et al., 2018). Spines change in size, shape and number depending on synaptic activity (for reviews see Bhatt et al., 2009; Harris, 2020; Kasai et al., 2010; Nishiyama & Yasuda, 2015; Segal, 2017; Suratkal et al., 2021), which has been termed extrinsic spine size dynamics (Kasai et al., 2021).

Given the overwhelming evidence for activity-dependent, extrinsic spine dynamics, the conventional view would be to expect spine size distributions to depend heavily on synaptic activity and associated synaptic plasticity (Barbour et al., 2007, see also Mateos et al., 2007; McKinney, 2010; McKinney et al., 1999). However, spines also display spontaneous, activity-independent, intrinsic changes (Mongillo et al., 2017; Ziv & Brenner, 2018). In keeping with a major role of such intrinsic spine dynamics, recent data from pharmacologically silenced cultured rat cortical neurons challenged the conventional view, indicating that skewed synapse weight distributions can emerge in an activity-independent manner (Hazan & Ziv, 2020). However, what remains unclear are the important questions as to (i) what kind of spine size distributions emerge during dendritic maturation of adult newborn neurons, when, and whether these are affected by homo- and heterosynaptic plasticity, and (ii) whether such skewed synapse weight distributions can emerge spontaneously in intact neuronal circuits. To address these issues, we studied the distribution of spine sizes in adult-born dentate granule cells from rats with induced *in vivo* homo- and heterosynaptic long-term plasticity. In addition, we studied spine size distribution in Munc13 double-knockout mouse brain circuits with completely blocked presynaptic activity. We found that homosynaptic long-term potentiation (LTP), with associated spine growth, and heterosynaptic long-term depression (LTD), with associated spine shrinkage, do not disrupt the lognormal-like spine size distribution but rather modulate its parameters. Moreover, we report that the lognormal-like distribution of spine sizes emerges even with entirely blocked synaptic activity.

## Results

### Independence of spine size distribution from long-term homo- and heterosynaptic plasticity in adult-born hippocampal granule cells (abGCs)

As the effects of nerve cell age and long-term synaptic plasticity on the skewness of spine size distributions are unknown, we characterized the spine size distribution and its relationship to long-term synaptic plasticity in retrovirally labeled hippocampal abGCs of three different cell ages. These are characterized by gradual onset and development of homo- and heterosynaptic plasticity (21, 28 and 35 dpi, see Methods; Jungenitz et al., 2018), soon after start of spinogenesis at 16-18 dpi (Ohkawa et al., 2012; Radic et al., 2017). In these cells, homosynaptic LTP associated with spine enlargement was induced in the middle molecular layer (MML) following 2 h stimulation of the medial perforant path (Jungenitz et al., 2018). At the same time, concurrent heterosynaptic LTD associated with spine shrinkage was induced in dendrites in the adjacent unstimulated outer and inner molecular layers (OML, IML). Those effects were restricted to the stimulated ipsilateral hemisphere and therefor, the unstimulated contralateral site served as control. Here, we fitted a lognormal function to the raw data to test whether it provides a good fit for the size distribution of mushroom spines. In the first round of analyses, this was done collectively for all of the cells of one condition (i.e. synaptic layer, cell age and hemisphere) together (Figure 1).

**Figure 1.**
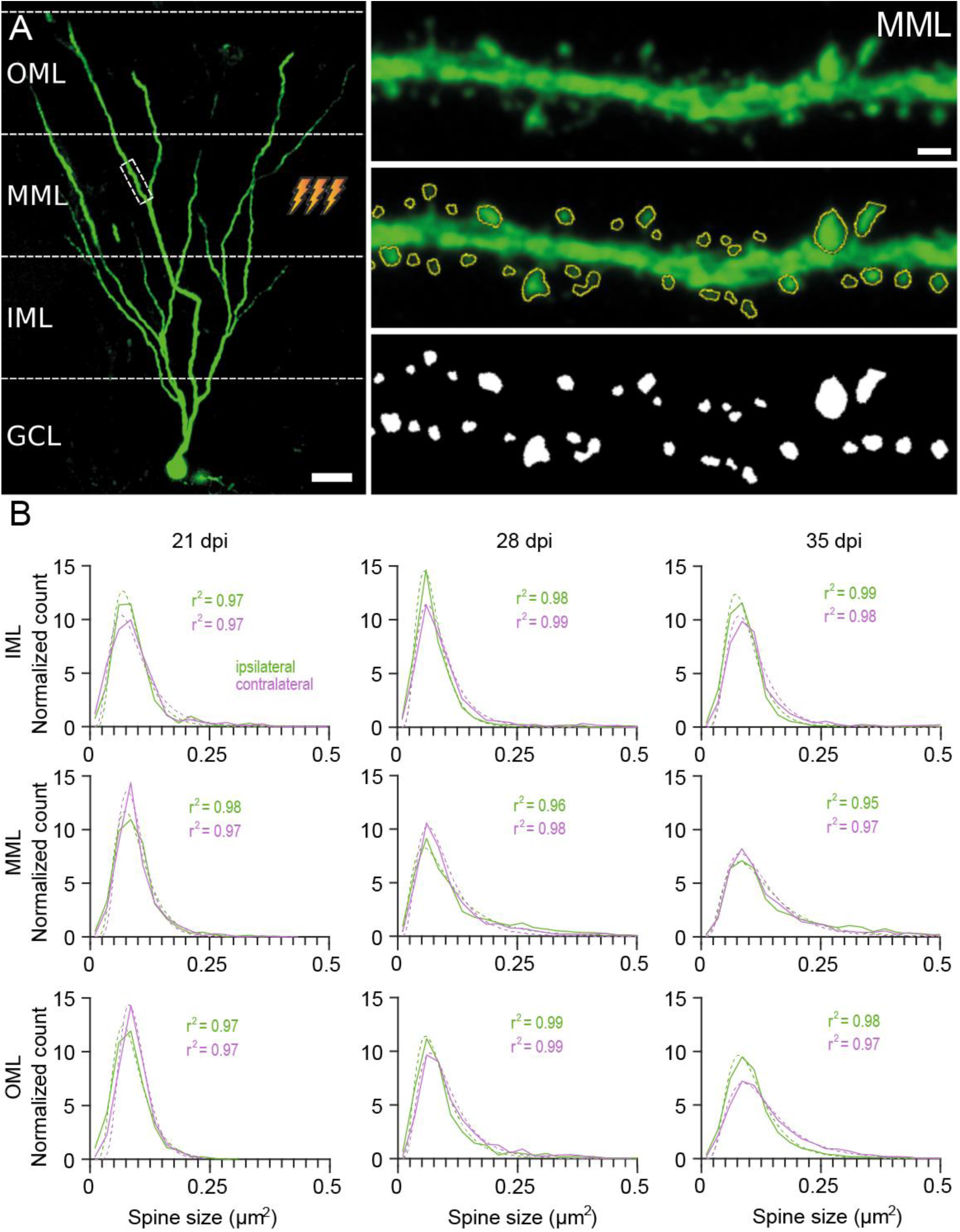
Collected spine size data from anesthetised rat abGCs reveal robust lognormal-like spine size distributions in all dentate layers and cell ages irrespective of ipsilaterally induced homosynaptic or heterosynaptic plasticity. (A) Left: An example retrovirally labeled abGC imaged at 35 days post injection (dpi; scale bar: 25 µm). The ipsilateral MML experienced 2-hour high frequency stimulation. Right: Top panel shows an enlarged dendritic segment located in the stimulated ipsilateral MML. Middle, bottom panel depicts analysed spines (scale bar: 1 µm). (B) Spine size distributions and their average lognormal fits for all cells in one layer (OML, MML, IML), time (21, 28 and 35 dpi = cell age) and hemisphere (ipsilateral stimulated = green, and contralateral control = magenta), fitted to the spine data. Note the high overall goodness of fit for all conditions. The lower ipsilateral vs. contralateral (stimulated vs. control) distribution peak associated with reduced distribution width in the stimulated MML indicates homosynaptic spine expansion; the higher ipsilateral vs. contralateral distribution peak in the OML and IML indicates heterosynaptic spine shrinkage. OML, MML, IML: outer, middle, inner molecular layer; GCL: granule cell layer of the dentate gyrus. The dashed line represents the lognormal fit, the solid line the spine data binned into size categories.

In all conditions – in ipsi- and contralateral dentate gyrus, at all cell ages and in all three layers – the lognormal-like distribution matched the data exceptionally well with very high goodness of fit (r^2^) values of 0.95 - 0.99. As expected, changes in the shape (peak and width) of the distribution reflected the overall homosynaptic spine enlargement in the ipsilateral MML with respect to the contralateral MML as well as the overall heterosynaptic spine shrinkage in the ipsilateral OML and IML with respect to the contralateral OML and IML. This confirms that after plasticity induction, the number of large spines increased and the number of small spines decreased in the stimulated layer while opposite changes occurred in the adjacent unstimulated layers (Jungenitz et al., 2018). However, the lognormal form of the distribution remained.

To see if a skewed, lognormal-like distribution also appeared at the level of individual cells, we examined spines in each cell separately. Both ipsilateral and contralateral (Supplementary Figure 1 and 2) dentate abGCs showed highly rightward skewed distributions at all cell ages and in all layers with a variety in shapes, peaks, and widths, and a lognormal-like spine size distribution was observed in all individual cells.

Overall, we achieved a good fit, with the majority of r^2^ values between 0.8 and 0.99. There was some variability in the goodness of fit as fewer samples were available for analysis, and one outlier was as low as -0.5 (MML ipsilateral, at 21dpi). The generally high r^2^ values indicate a lognormality of the data at the individual cell level, independent of cell age, cell layer, or stimulation (hemisphere). Thus, the rightward skewness of spine size distribution is a robust and synaptic-plasticity-independent phenomenon already present at an early nerve cell age.

To quantify the comparison of spine size distributions between ipsilateral (stimulated) dentate gyrus with induced synaptic plasticity and the contralateral (control) side, we calculated the goodness of fit (r^2^) (Figure 2) and skewness (Supplementary Figure 3).

**Figure 2.**
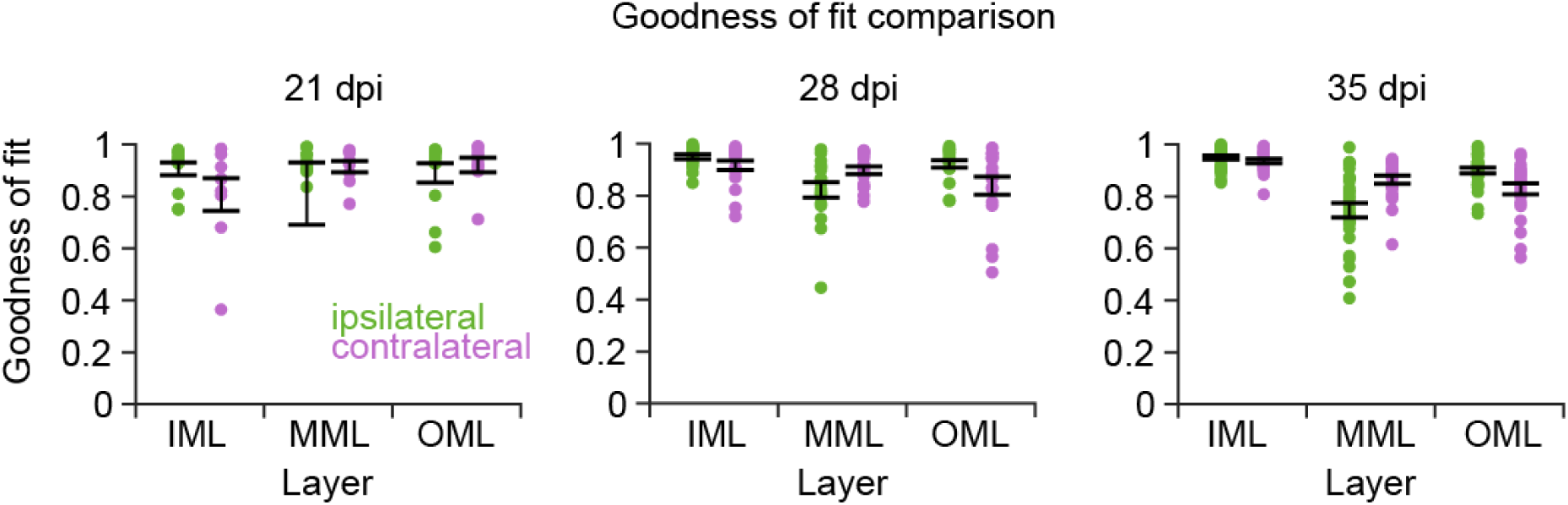
Individual cell level analysis of the spine size data from anesthetised rat abGCs confirms robust lognormal-like distribution in all dentate layers and cell ages. The goodness of fit values were similar in the ipsilateral (stimulated; green) and contralateral (control; magenta) dentate gyrus layers. Left, middle, right panel: 21, 28 and 35 dpi, respectively. Left, middle, right panel: 21, 28 and 35 dpi, respectively. Each dot represents a single cell. The error bar represents SEM. The y-axes are cropped at 0, with one outlier below this value in the MML ipsilaterally at 21 dpi.

There were no significant differences (p < 0.05) in the goodness of fit between the two hemispheres, in any layer or time (i.e. cell age in dpi) comparison (Figure 2). Another way to quantify the lognormality of the spine size data is to calculate the skewness (asymmetry around the mean) of the data. All cells in every condition displayed a skewness above 0, confirming that the data were not symmetrically distributed but skewed to the right (Supplementary Figure 3). Again, there were no significant differences between the hemispheres. Overall, the skewness quantification supported the results obtained by the r^2^ comparisons, showing that the lognormal-like distribution of spine sizes is independent of stimulation-induced homo- and heterosynaptic plasticity. Comparing the standard deviations taken from the natural logarithms of the spine data (in the following called sigma), which is an indicator of the width of the distribution and in this case the range of the spine sizes, some significant differences (p < 0.05) were observed (Supplementary Figure 4). The sigma value for the stimulated ipsilateral MML at 28 dpi significantly increased compared to the contralateral side. This indicates that the shape widened and that there was an increase in bigger spines due to the induction of homosynaptic LTP. There was a significant decrease in the ipsilateral spine sizes in the IML at 21 dpi and the OML at 35 dpi compared to the contralateral side, indicating that the shape narrowed and the number of smaller spines increased due to heterosynaptic LTD.

For a lognormal distribution, the logarithm of the individual values is normally distributed. As an additional quantification method, we calculated the logarithm of the data and fitted a Gaussian distribution to the transformed data (Figure 3). The distributions at the youngest cell age (21 dpi) showed a well-fitted Gaussian distribution in all three layers and both ipsi- and contralaterally, indicating the condition for the lognormal distribution was met. In older cells (both 28 and 35 dpi), the Gaussian distribution fit less well to the logarithmic data. This was especially the case on the right side of the peak, where the actual number of spine sizes was higher than the estimated fit. There was an overabundance of bigger spines at older cell ages, regardless of plasticity induction. However, this overabundance of bigger spines could be observed especially in the MML, where homosynaptic plasticity was induced. This indicates that spines do not follow a strict lognormal distribution but a lognormal-like distribution.

**Figure 3.**
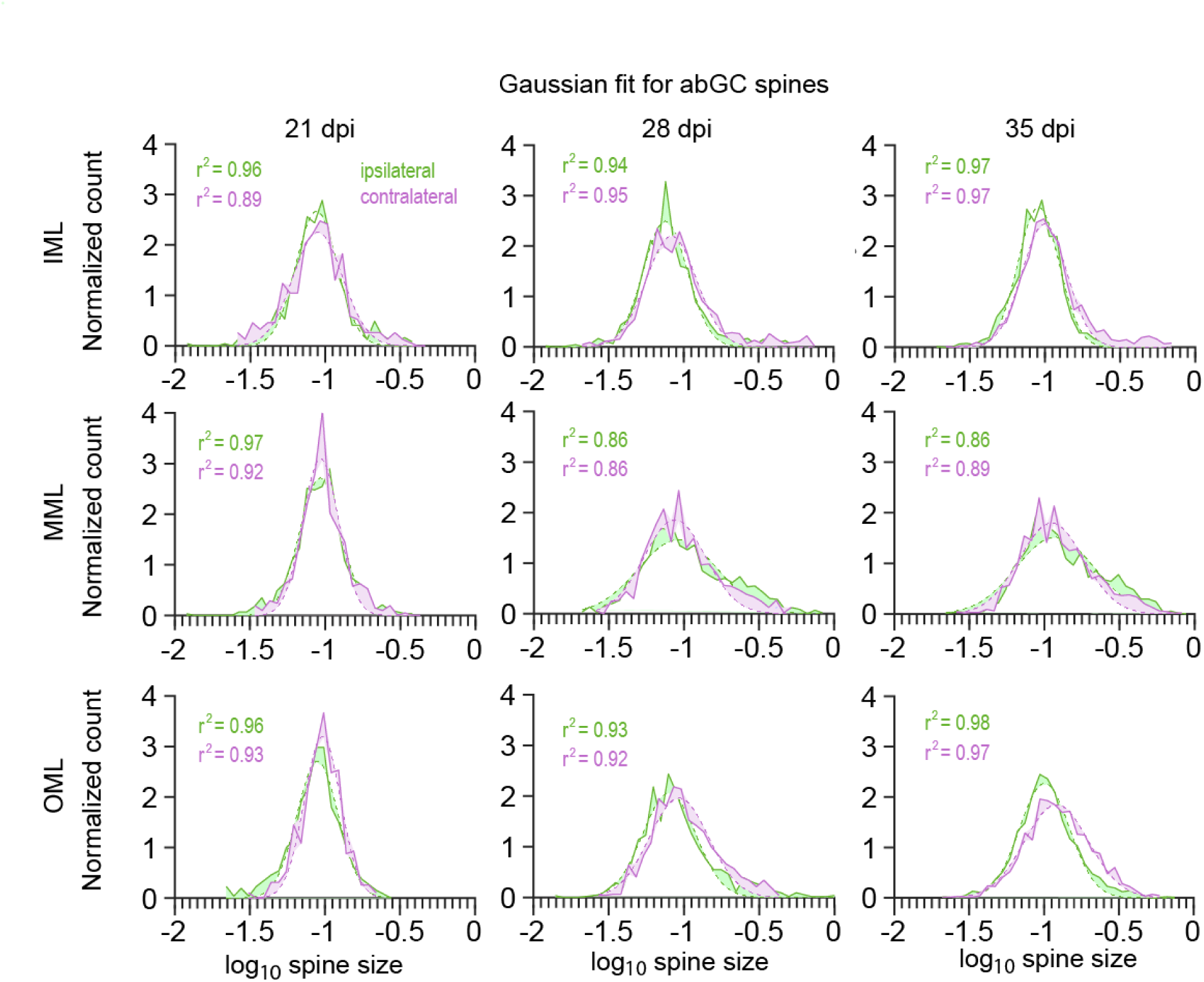
Fitting a Gaussian distribution to logarithmically transformed spine data of abGCs revealed that spine sizes follow a lognormal-like distribution. The average Gaussian fits for all cells in one dentate layer (OML, MML, IML), time (21, 28 and 35 dpi) and hemisphere (ipsi- and contralateral), fitted to the logarithm of the spine data (green and magenta). The dashed line shows the Gaussian fit, the solid line represents the spine data. The differences between data and fit is shown by shading in the areas between both. At 21 dpi, in all layers and both ipsi- and contralateral, the Gaussian distribution fits well to the data. In the older cells (28 and 35 dpi) there is apparent overabundance of bigger spines and thus a bias to the right of the peak. This is pronounced especially in the MML, where high-frequency stimulation occurred. OML, MML, IML: outer, middle, inner molecular layer; GCL: granule cell layer of the dentate gyrus.

We compared three skewed distributions, including the lognormal distribution, to quantify whether the lognormal distribution is the best fit of those three. To this end, we used the Akaike Information Criterion (AIC). Our analyses and comparisons revealed that of the three distributions tested (lognormal, gamma and Weibull), the lognormal distribution had an advantage over the other two, indicating that it was the best fit for the data (Supplementary Figure 5 and 6).

### Independence of spine size distribution from presynaptic transmitter release

Viewed together, the data from rat abGCs showed a strong independence of the lognormal-like spine size distribution from homosynaptic and heterosynaptic plasticity. This raises the question as to whether synaptic activity in general affects spine size distributions. To assess this, we analysed spines in nerve cells with blocked presynaptic transmitter release.

We used a data set of CA1 pyramidal cell (CA1 PC) spines from organotypic hippocampal cultures obtained from Munc-13-1-Munc13-2 double knockout mice (DKOs) (Sigler et al. 2017). In these mutants, presynaptic glutamate and GABA release is almost entirely blocked (Varoqueaux et al. 2002; Sigler et al. 2017). The spine data comprised three developmental time points, at which spine size was measured in organotypic slices (7, 14 and 21 days in vitro, div) and two further groups, one where synaptic activity (presynaptic transmitter release) was blocked (DKO group 0) and the corresponding control group (group 1). CA1 PCs possess three different spine types: 22.85±6.01% mushroom spines (mean±SD), 23.73±4.83% thin spines and 51.16±6.62% stubby spines. About 2.26±2.53% were defined as ‘other’ and not included in further analyses.

The data were analysed by different conditions, separated by time *in vitro* (div) and group. In the first step, all cells and spine types were analysed together in each condition. In the second step, spine sizes were analysed at the single cell level, for all spine types together. Finally, the three different spine types were analysed separately, first for all cells in one condition, then at the individual cell level as well.

A lognormal distribution was fitted to the spine data. As with the abGC data above, the goodness of fit (r^2^) showed that the lognormal fit described the spine size distribution very well, in all conditions and for all spines (Figure 4).

**Figure 4.**
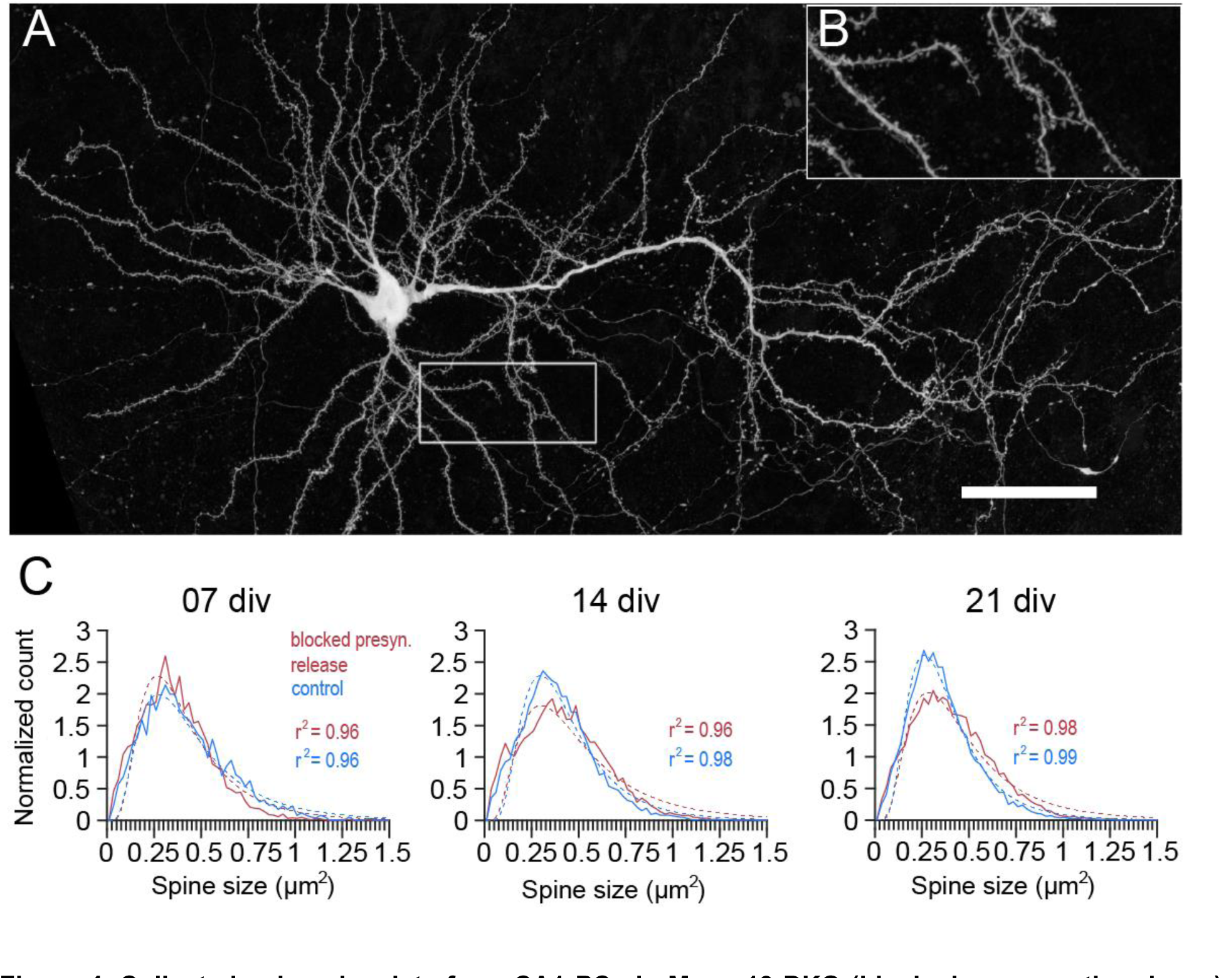
Collected spine size data from CA1 PCs in Munc-13 DKO (blocked presynaptic release) and WT (control) organotypic slice cultures revealed robust lognormal-like distribution in all cell culture ages irrespective of blocked presynaptic release. (A) An example GFP labeled CA1 PC from a DKO slice culture imaged at 21 days in vitro (div; scale: 50 µm). (B) The panel shows an enlarged dendritic segment. (C) Average lognormal fit for all spines (mushroom, stubby and thin) and all CA1 PCs in one condition (blocked presynaptic release or control) pooled together. Note that the lognormal function fit the data (red and blue dots) with high goodness of fit (r^2^) values in both groups, and at all time points (div). The dashed line shows the lognormal fit, the solid line represents the spine data.

Again, like with the abGC data, at the individual cell level, spine sizes in every CA1 PC in both groups followed a lognormal distribution, at each cell culture age (div) that we studied (Figure 5A). There were differences in the shape and width of the distribution, but the rightward skewness was preserved even at the individual cell level.

**Figure 5.**
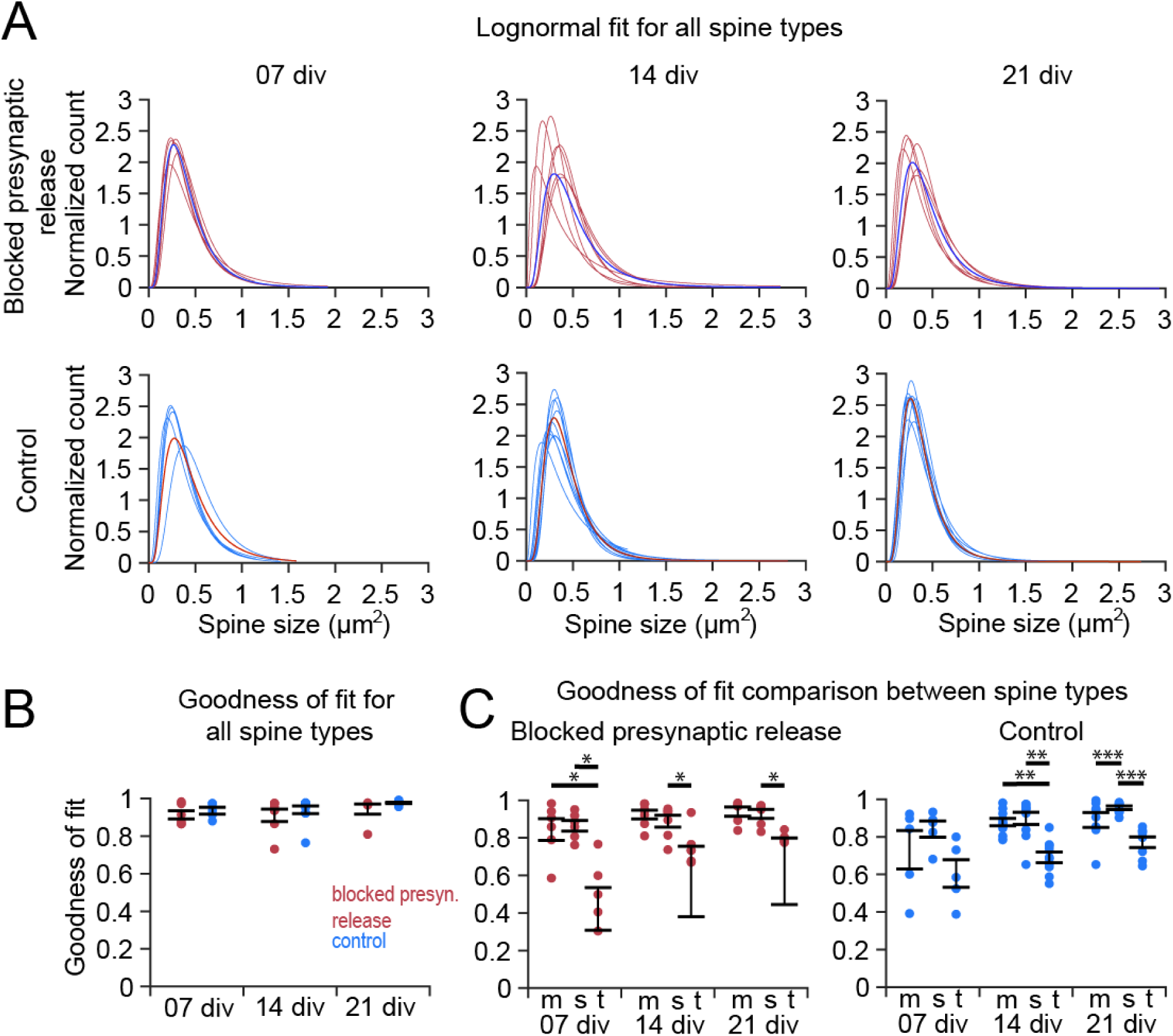
Individual cell level analysis of the spine size data from CA1 PCs in Munc-13 DKO (blocked presynaptic release) and WT (control) organotypic slice cultures revealed a robust lognormal-like distribution independent of synaptic activity. (A) Lognormal fits in individual cells in both groups and at three time points (div). The single blue (above) and red (below) line represents the mean of all spine sizes as seen in Figure 4C. (B) Goodness of fit (r^2^) comparisons. The comparison between the two groups yielded no significant difference. In the control group, r^2^ increases significantly (p < 0.01) over time. (C) Thin spines showed lower goodness of fit than stubby and mushroom spines in both experimental conditions. Each dot represents a single cell, error bars represent SEM. m – mushroom spines, s – stubby spines, t – thin spines

We compared the goodness of fit parameter r^2^ between the groups and different time points (div), for all spines together (Figure 5B). There were no significant differences between the two groups, only a trend in the blocked activity group towards a slightly reduced r^2^. Comparing the time points, there was no significant difference (p < 0.05) within the blocked activity group. In the control group there was a significant increase (p < 0.01) in the goodness of fit from day 7 to day 21 *in vitro*, indicating that the lognormal distribution described the data better for more mature slice cultures. A similar trend was seen in the blocked activity group, but without reaching statistical significance. This shows that there is a lognormal-like distribution of spine sizes irrespective of whether the presynaptic transmitter release is blocked or not.

A closer analysis of the spine size data revealed that the skewness values were typically above 0 (in some exceptional cases for thin spines below 0, indicating a skewness to the left), confirming that the spine sizes were not symmetrically distributed. Comparing the different conditions revealed no significant differences between different time points (cell culture age in div) or between the groups (Supplementary Figure 7A). Within each group, the skewness increased slightly but not significantly over time. The sigma comparison revealed no significant differences in the width of the distribution and the range of spine sizes (Supplementary Figure 8). A trend was seen at 21 div, where the blocked presynaptic transmitter release group has a slightly increased sigma compared to the control group, indicating that the range of spine sizes increases.

Next, we tested whether a deeper analysis of spine type subgroups (mushroom, stubby and thin) would show inter- or intra-group differences (Figure 5C). The thin spine population showed lower r^2^ values than the mushroom and stubby spine population. In line with this, thin spines also showed the lowest score for skewness (Supplementary Figure 5B). At the individual cell level, mushroom spines in each cell followed a lognormal distribution (Figure 6A). The group with blocked presynaptic transmitter release showed a similar goodness of fit as the control group. There was a significant increase of r^2^ over time (p < 0.05) in the control group (Figure 6B). Mushroom spines had a slightly higher skewness in the control group, but the difference was not significant (Supplementary Figure 7C) and they showed the lowest sigma value in comparison to the other two spine types, indicating a smaller range of sizes (Supplementary Figure 8). Analyses of thin and stubby spines at the individual cell level are shown in Supplementary Figures 9 and 10.

**Figure 6.**
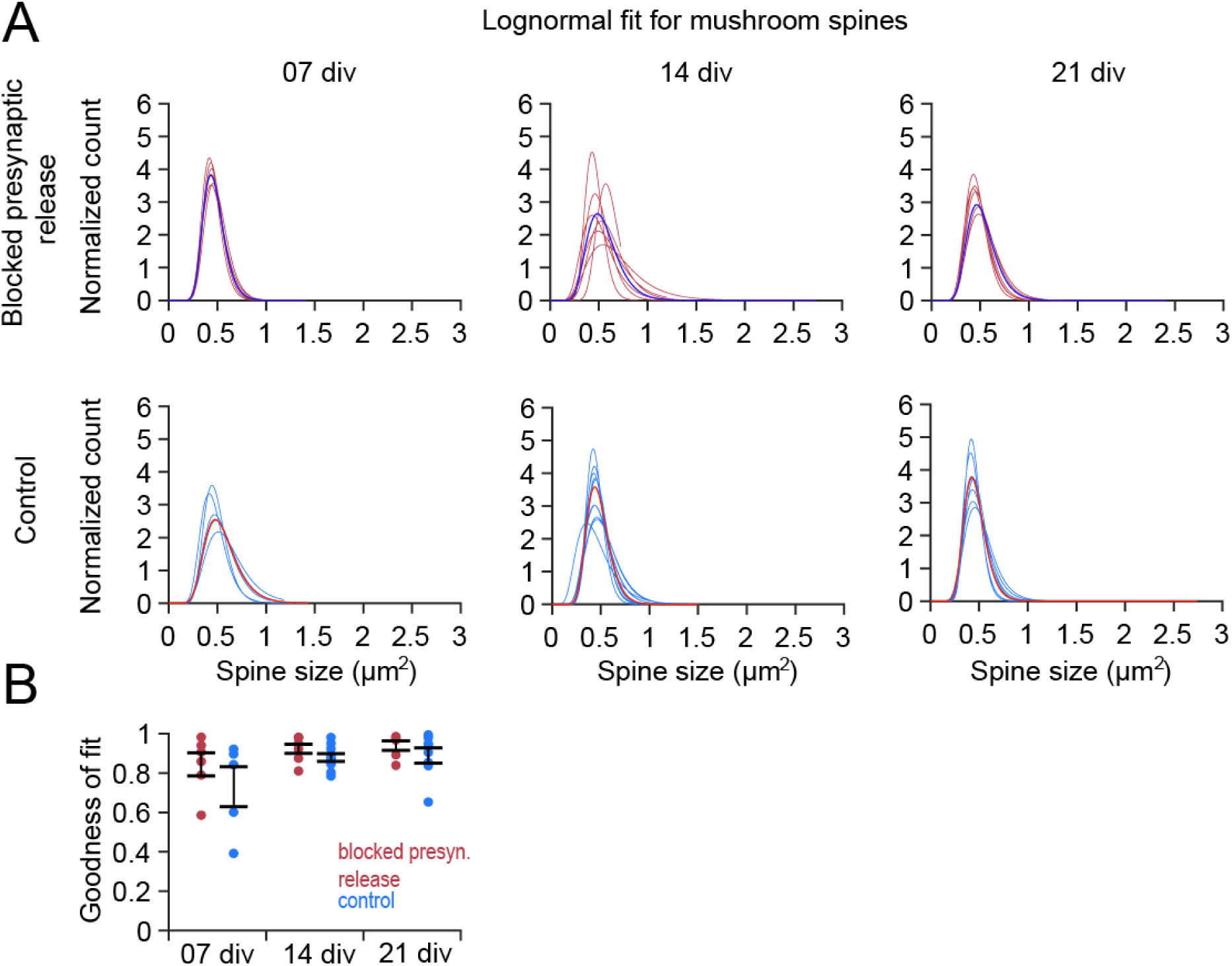
Analysis of mushroom spines from CA1 PCs in Munc-13 DKO and WT (control) organotypic slice cultures showed a high goodness of fit to a lognormal distribution. (A) Individual fits for mushroom spines in each cell. The single blue (above) and red (below) fit shows the average distribution (B) Goodness of fit (left panel) analysis revealed no significant differences between the groups (blocked presynaptic release vs. control) and a significant increase in r^2^ over time (p < 0.05) (7, 14 and 21 div) for control. Each dot represents a single cell, error bar represents SEM.

As with the abGC data set, we conducted the AIC analysis and comparison for mushroom spines, to check whether the lognormal distribution was the best fit out of three skewed distributions. The lognormal distribution had an advantage over the other two in both experimental groups and at all cell ages (Supplementary Figure 11). These findings indicate that a lognormal-like spine size distribution is preserved even when synaptic activity is blocked. Intriguingly, the sizes of thin spines showed a less good fit to a lognormal distribution.

Again, as with the abGC spine data set, a final analysis of spine data from Munc13 DKOs and control littermates focused on the lognormal-like distributions of spine sizes in more detail by employing the normal (Gaussian) fits of logarithmically transformed data. The logarithm of lognormal-like spine size data should lead to a normal-like distribution. Taking the logarithm of the data and fitting a Gaussian distribution to the transformed data revealed for all spine types that the distribution had a bias towards the left side of the peak, meaning there was an overabundance of small spines in the samples (Figure 7A), at all cell ages and in both experimental groups. For mushroom spines, there was a clear cutoff to the left (Figure 7B), whereas for thin spines there was a cutoff to the right of the peak (Figure 7C). This was due to the method by which spines were categorised by size into thin or mushroom spines. Stubby spines showed the best Gaussian fit, indicating that the stubby spines were distributed strictly lognormally. The bias to the left might be an artefact of the method used to detect and measure the spines. Overall, the findings indicate, similar to the abGC data set, that spines were lognormal-like distributed independently of synaptic activity.

**Figure 7.**
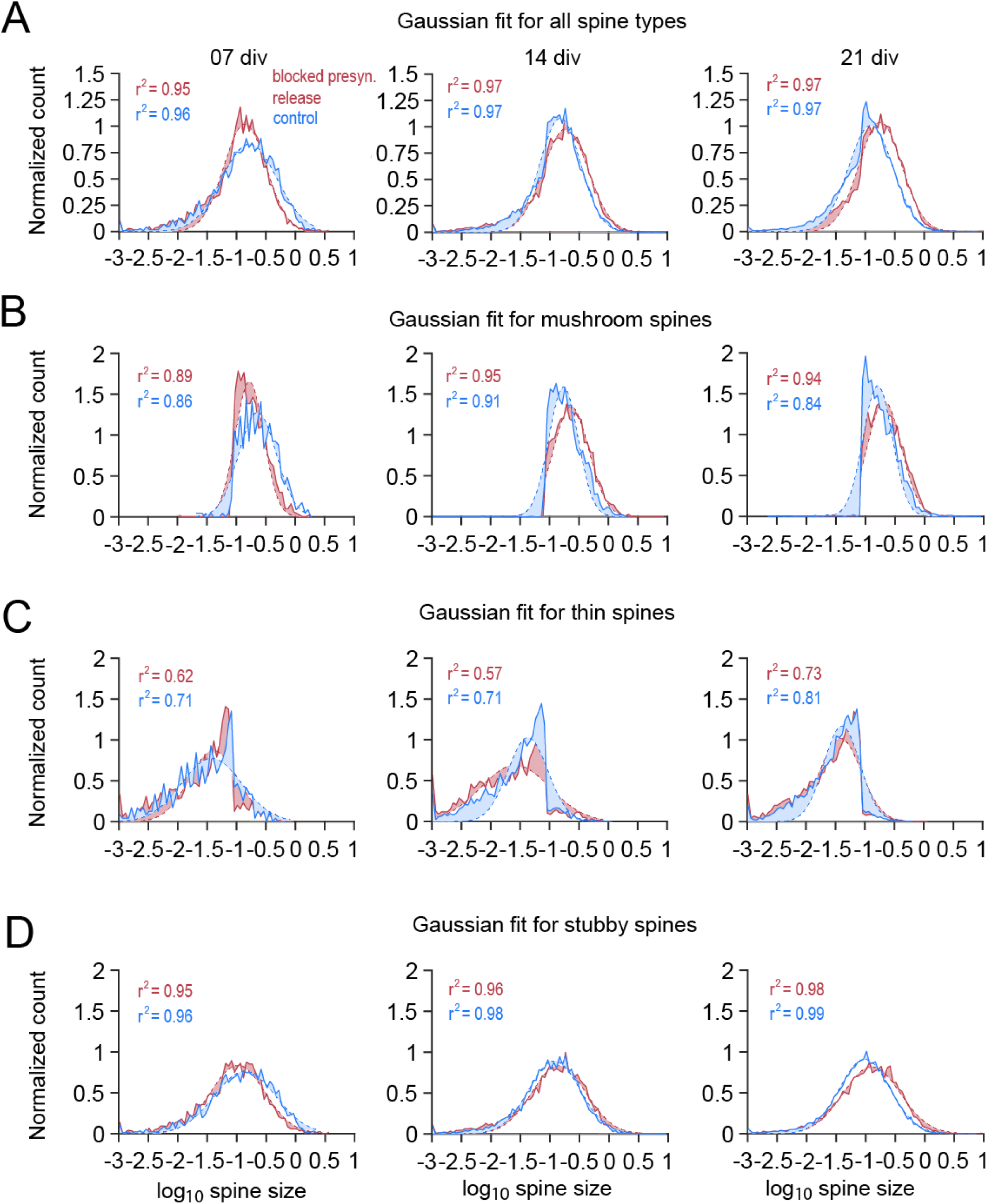
Gaussian fits to logarithmically transformed spine data for all spine subtypes and each type individually showed varying degrees of lognormality. The average Gaussian fits for all spine subtypes (A), mushroom spines (B), thin spines (C) and stubby spines (D) in all cells at one time (7, 14 and 21 div) and experimental condition (blocked presynaptic release and control), fitted to the logarithm of the spine data (red and blue). For all spine types together, there was a bias to the left of the peak at all three cell ages and for both conditions, indicating an overabundance of small spines in the data sample, making the distribution more lognormal-like. Mushroom spines (B) and thin spines (C) showed a cut off at the same spine size, with mushroom spines displaying spine sizes above the cutoff and thin spines below. The Gaussian distribution did not fit as well to those two spine types, meaning that they are more lognormal-like. Stubby spines (D) showed the best fit to the logarithmic data. The dashed line shows the Gaussian fit, the solid line represents the spine data. The differences between data and fit is shown by shading in the areas between both.

### A computational model implementing intrinsic and extrinsic synaptic plasticity accounts for the generation and preservation of skewed synaptic weight distributions

Many computational models of synaptic dynamics presume that the distribution of synaptic weights arises predominantly due to activity-dependent (extrinsic) synaptic plasticity (Gilson and Fukai, 2011; Zheng et al., 2013; Effenberger et al., 2015; Scheler, 2017). Therefore, our observation that synaptic activity is not necessary for the emergence of skewed spine size distributions requires an extended computational approach that captures the key role of activity-independent (intrinsic) plasticity. To account for this, we used a computational model of synaptic dynamics that combines intrinsic plasticity (Hazan and Ziv, 2020) with classical extrinsic plasticity mechanisms.

Lognormal distributions are typically preserved when applying multiplicative stochastic operations. Combined intrinsic and extrinsic synaptic plasticity might represent a biological implementation of such multiplicative changes of synaptic weights. Thus, to investigate the influence of intrinsic and extrinsic plasticity on the lognormal distribution of spine sizes, we developed a minimal computational model, that was able to account for the experimental data. Extrinsic synaptic plasticity was modeled as Hebbian activity-dependent spike-timing-dependent plasticity (STDP) consisting of additive LTP and multiplicative LTD. Intrinsic synaptic plasticity was based on activity-independent fluctuations modeled as multiplicative noise. The model was inspired by van Rossum et al. (2000). The synaptic weights, for which we assume spine sizes to be a reliable proxy, were determined for each condition after the simulation was run, and a lognormal distribution was fitted over the weight data. In a first simulation, we wanted to see if intrinsic plasticity alone (modeled as multiplicative noise) can generate a lognormal distribution. To this end, we fed a uniform distribution as initial weights into the model and tracked the synaptic weights over the time course of the simulation to see how it developed (Figure 8A). The distribution became lognormal over time, showing that multiplicative noise is indeed sufficient to generate lognormal distributions (Hazan & Ziv, 2000).

**Figure 8.**
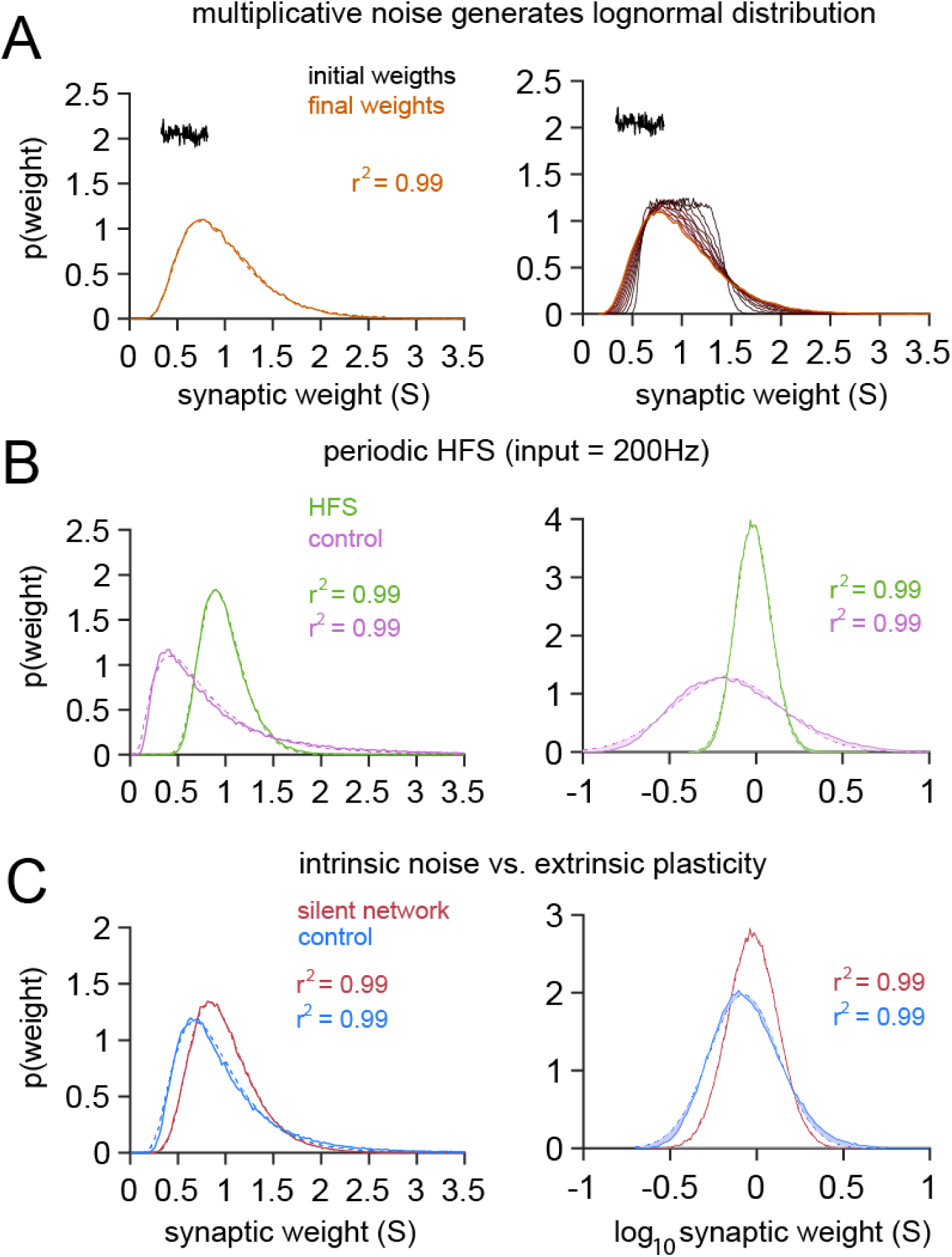
Computational modeling indicates that intrinsic (activity-independent) synaptic plasticity is sufficient for the generation of lognormal-like spine size distributions and a combination of intrinsic and extrinsic synaptic plasticity is sufficient for the maintainance of the lognormal-like distributions. (A) An initially uniform distribution transforms into a lognormal distribution over time. Only multiplicative noise that represented intrinsic synaptic plasticity was applied to the distribution, which was enough to generate a lognormal distribution over time out of the uniform distribution. (B) Periodic high frequency simulation (HFS) with an input rate of 200 Hz was applied to the model (light green) and compared to a control simulation that received a 10 Hz periodic input (magenta). The HFS simulation shows a narrower distribution with the peak centered around medium sized weights whereas the control simulation has a broader distribution with the peak more to the left, centered around smaller weights. Both simulations follow a lognormal distribution with high goodness of fit values. (C) Intrinsic mechanisms, simulated as a silent network without any extrinsic input (red) shows a slightly shifted distribution compared to the control simulation (extrinsic activity simulated with an input at 5 Hz; light blue). The silent simulation has a slightly higher peak centered around medium sized weights and seems to be broader compared to the control simulation. Both simulations follow a lognormal distribution with high goodness of fit values.

Next, we explored *in silico* how synaptic activity in the form of LTP-inducing HFS affects the shape of the synaptic weight distribution. We used the model to recreate the plasticity processes in the Jungenitz et al. (2018) data set. We compared a HFS (periodic spiking input at 200 Hz) with a control simulation with an input of 10 Hz (Figure 8B). The model generated lognormal distributions in both simulations with high goodness of fit values (r^2^ = 0.99), but with differences in shape and peak. This was supported by the Gaussian distribution of the logarithmic weight data (Figure 8B, right panel). The lognormal distribution resulting from the HFS simulation showed a narrower distribution and a higher peak at medium sized spine sizes, whereas the control simulation showed a broader shape, with a peak at lower sized weights. This is contrary to the experimental results, where the HFS stimulated spines in the MML showed a broader distribution with bigger spines increasing in number, whereas the unstimulated control spines showed a narrower distribution. This discrepancy could be due to the differences in the duration of the experiment and the simulation, or to the choice of modelled heterosynaptic scaling.

We then recreated the experimental data obtained upon block of presynaptic transmitter release by comparing a completely silent simulation (i.e. using only intrinsic noise) and a control simulation that received Poisson input at a frequency of 5 Hz (Figure 8C). The simulation yielded similar results to the experimental data. The intrinsic noise distribution broadened as compared to the control simulation, but there were no significant differences between the two groups. Both distributions were well-fitted by a lognormal distribution (r^2^ = 0.99), which was also supported by the well-fitted Gaussian distribution of the logarithmic weight data (Figure 8C, right panel). As shown by Rossum et al (2000), additive STDP also contributes to a skewed distribution of synaptic weights as already strong synapses are more likely to trigger a postsynaptic response and therefore potentiate again. Interestingly, however, additive intrinsic noise can lead to relatively large changes in the strengths of small synapses and limit the skewness of the weight distributions. This, alongside the experimental results on silenced cultures, implies that intrinsic noise should be chiefly multiplicative.

In sum, and in agreement with the abGC spine data, combined extrinsic and intrinsic plasticity can maintain the skewed distributions in the presence of correlated LTP-inducing synaptic activation. Furthermore, in line with Munc-13 DKO spine size data, our modeling shows that extrinsic plasticity is not necessary for the generation of skewed spine size distributions and that intrinsic plasticity alone is sufficient.

## Discussion

Excitatory post-synaptic potential sizes and spine head sizes have lognormal-like distributions (de Vivo et al., 2017; Ikegaya et al., 2013; Lefort et al., 2009; Loewenstein et al., 2011; Merchán-Pérez et al., 2014; Montero-Crespo et al., 2020; Santuy et al., 2018; Song et al., 2005). Here, we confirm that spine size distributions follow a lognormal shape in both hippocampal dentate abGCs *in vivo* and in organotypically cultured CA1 PCs. In dentate abGCs, a lognormal-like distribution of spine sizes was present at all studied cell ages, irrespective of homo- or heterosynaptic long-term plasticity induction. Most strikingly, in CA1 PCs, spine size distributions were skewed and lognormal-like even in Munc-13 DKOs, in which presynaptic transmitter release is entirely blocked. These data show that the lognormal-like distribution of spine sizes is activity- and plasticity-independent. The skewness of spine size distributions develops early in cell age without extrinsic influences related to presynaptic transmitter release, and therefore seems to be determined intrinsically. However, we cannot exclude potential extrinsic influences that are not related to presynaptic transmitter release, such as trophic factors or adhesion proteins.

### Independence of spine size distributions from intrinsic dynamics and extrinsic plasticity

Intriguingly, we detected robust lognormal-like distributions of spine sizes in young newborn GCs that had experienced homo- and heterosynaptic plasticity. This is in agreement with previous studies showing unchanged spine size, spine type distribution, and spine numbers at 30 min and 2 h after homosynaptic long-term potentiation in dentate granule cells and CA1 PCs, respectively (Bromer et al., 2018; Sorra & Harris, 1998). Together with our previous work (Jungenitz et al. 2018, Beining et al., 2017), these data indicate that high-frequency activation of synapses evokes their homo- and heterosynaptic plastic changes leading to a redistribution instead of an overall increase (or decrease) in spine size and synaptic strength. The plasticity-related redistribution of synaptic weights with a homeostatic maintenance of the total synaptic area per µm of dendrite length (Bourne & Harris, 2007; Bromer et al., 2018) may be a result of activity-dependent competitive redistribution of synaptic building resources (Triesch et al., 2018). In addition to the plasticity-independence, the skewed spine size distribution in abGCs was detected at the earliest studied time point (21 dpi), shortly after onset of spinogenesis between 16 – 18 dpi (Ohkawa et al., 2012; Radic et al., 2017). This indicates that it develops in early stages of a nerve cell’s life. Extending long-term time lapse imaging of abGCs (Radic et al., 2017) to include their initial developmental stages with the time of rapid spinogenesis should clarify whether the first spines already display skewed size distributions.

A recent study on cultured primary cortical neurons (Hazan & Ziv, 2020) provided results in line with our observation that spine size distribution is independent of presynaptic glutamate release. In this study on dissociated neurons in culture with pharmacologically blocked spiking and synaptic activity during the plating procedure, synapses showed physiological diversity with a full range of synaptic sizes (Hazan & Ziv, 2020). The synapse size distributions in these silenced networks in culture were rightward skewed, broad, and stable, showing characteristics of a lognormal-like distribution. Interestingly, networks with chronic activity suppression showed an increase in average spine size, and synaptic size distributions broadened, indicating that activity-dependent processes constrain synaptic growth (Minerbi et al., 2009; Statman et al., 2014; Ziv & Brenner, 2018).

Our analysis of spines upon blockage of presynaptic transmitter release documents a similar shift in spine sizes. The blocked transmitter release group shows a broader distribution with a lower peak, indicating a shift towards an increased number of bigger spines, possibly regulated by intrinsic mechanisms. Similar results were reported by Yasumatsu and colleagues (2008) who observed individual spines of CA1 pyramidal cells from rat hippocampal slices in culture after blocking synaptic transmission and plasticity mediated by NMDA receptors. They reported that spontaneous, intrinsic spine volume fluctuations were independent of activity-dependent plasticity processes. In the presence of NMDAR inhibition, the rate at which spines were eliminated was decreased and spine generation was unaffected. Spine elimination of mostly small spines was reduced but new, small spines still emerged, affecting the skewness of the distribution.

An important finding of Yasumatsu et al. (2008) was that small spines were the most plastic ones, changing in size, being eliminated, or newly generated even within one day. Large spines, in contrast, were more persistent. This supports the idea that small, more plastic spines are more involved in learning processes, whereas stable, large spines are responsible for memory traces (Bourne & Harris, 2007; Hung et al., 2008; Kasai et al., 2003). This might hint at a potential advantage of lognormal size distributions, with a large pool of small spines with higher plasticity potential and a minority of big and less plastic spines that can hold long-term memory traces (cf. Yap et al., 2020). However, our present study and previously published data (Hazan & Ziv, 2020; Sando et al., 2017; Sigler et al., 2017, Kleinjan et al., 2023) show clearly that synaptic activity is not necessary for the emergence of large spines (Ziv & Brenner, 2018). In line with this, the diversity of spine types – in terms of fractions of mushroom, stubby and thin spines – is not affected in mice with a complete suppression of synaptic transmitter release from glutamatergic neurons upon Cre-inducible expression of tetanus toxin (Dorkenwald et al., 2019; Sando et al., 2017). Consistently, spinogenesis in CA1 PCs has been shown to be independent of the activation of ionotropic glutamate receptors (Lu et al., 2013), although their numbers might be modulated by the lack of activity (Sigler et al, 2017; Hazan and Ziv, 2020). Even the complete knockout of Ca^2+^ channels in synapses in cultured hippocampal neurons did not impair synapse structure (Held et al., 2020). All these observations are congruent with early investigations showing that *in vivo*-like synapse diversity emerges in neurons in chronically silenced organotypic cultures (Harms et al., 2005; Harms & Craig, 2005; van Huizen et al., 1985; but see McKinney et al., 1999).

### Computational model accounts for the generation and maintenance of lognormal-like weight distributions

The finding that synaptic activity is not necessary for the skewed spine size and synapse weight distribution is unexpected in the context of several prominent theoretical models. Many computational models of synaptic weight dynamics assume that realistic weight distributions emerge due to a combination of Hebbian and non-Hebbian activity-dependent synaptic plasticity. For example, spiking network simulations led to the suggestion that a highly skewed distribution of synaptic weights appears due to network self-organization (Zheng et al., 2013), by the combined effects of (i) excitatory and (ii) inhibitory spike-timing dependent plasticity (STDP and iSTDP), (iii) synaptic normalisation (preserving the total input weight of a neuron), (iv) intrinsic plasticity (for firing rate homeostasis), and (v) structural plasticity (in the form of synaptogenesis).

Similarly, other computational studies (van Rossum et al., 2000) used an STDP rule with a homeostatic component (diminished potentiation for strengthened synapses; see also Effenberger et al., 2015) or log-STDP (Gilson & Fukai, 2011) to reproduce the experimentally observed positively skewed weight distribution. Further, a more recent mathematical study argued that Hebbian learning is needed to produce and maintain skewed synapse size distributions (Scheler, 2017). However, the studies including our work and work of others (Hazan & Ziv, 2020) clearly show that activity-dependent synaptic plasticity is not essential for the lognormal-like weight distributions to occur. This means that the synaptic plasticity rules proposed in these computational studies are not necessary for the generation of heavy-tailed synaptic weight distributions, but that they may still be involved in the maintenance of the skewed distributions once neuronal networks become exposed to prolonged synaptic activity and plasticity.

Indeed, our plasticity model, using a Kesten process as multiplicative noise for implementing intrinsic synaptic fluctuations (Hazan & Ziv, 2020), generated a lognormal-like distribution without any influence of an extrinsic plasticity mechanism. The multiplicative noise (i.e. intrinsic plasticity mechanisms) also generated a lognormal distribution that is slightly broader than a control simulation with noise and activity-dependent plasticity (i.e. both intrinsic and extrinsic mechanisms). This is in accordance with our results obtained with the Munc13 DKO data set. When we added additive STDP and simulated the network model with periodic high-frequency input (mimicking LTP-inducing activity), the skewed, lognormal-like distribution was maintained, but changed in width and shape compared to a control simulation that received 10 Hz input. The maintenance of the lognormal-like distribution is in agreement with the abGC LTP/LTD data set.

However, in experimental data, the spine distribution affected by LTP-inducing HFS broadened compared to the control spines. In our model, the weight distribution of the HFS simulation got narrower in comparison to the control simulation. This discrepancy between the model and the data could be due to the time differences between the experiment and the simulation, where the HFS was applied for 2 hours to the cells, compared to the shorter period of the computer simulation. It could also be due to the choice of heterosynaptic scaling in the model. Further work is required to establish the range of parameters that are fully consistent with the experimental data, for example the relationship between strengths of intrinsic and extrinsic plasticity. The insight of this model, as previously shown by van Rossum (2000), is that even additive potentiation can generate and preserve skewed synaptic weight distributions as stronger synapses are more likely to trigger postsynaptic spikes and therefore more likely to undergo potentiation. The presence of skewed distributions even without STDP in our data is evidence that intrinsic noise is likely to be multiplicative.

Activity-independent computational models based on stochastic multiplicative shrinkage and additive growth of synapses (mathematically well approximated by stochastic Kesten or nonlinear Langevin processes) successfully account for the emergence of lognormal-like synaptic strength distributions (Hazan and Ziv 2020; see also Yasumatsu et al. 2008 and Loewenstein et al. 2011). Similarly, a mechanistic model based on activity-independent cooperative stochastic binding and unbinding of synaptic scaffold molecules can explain the rightward skewed, distributions of synaptic sizes (Hazan & Ziv, 2020; Shomar et al., 2017). Our new model of intrinsic and extrinsic plasticity shows how activity-independent and activity-dependent synaptic dynamics may cooperate to maintain lognormal-like distribution of synaptic efficacies.

An open question that remains is as to whether long-tailed distributions of synaptic weights have functional relevance. Their computational role is still not fully understood but several studies indicate that they may support optimal network dynamics in the form of sparse, fast, broad and stable responses (Cossell et al., 2015; Ikegaya et al., 2013; Iyer et al., 2013; Teramae et al., 2012; Teramae & Fukai, 2014) and facilitate network burst propagation (Omura et al., 2015). Sparse and strong synapses connect together to a so-called “rich club” of rare but highly connected neurons (Gal et al., 2017; Nigam et al., 2016). The rich-club neuron organization can generate bistable low-firing and high-firing network states, whereas biologically unrealistic random networks only display mono-stable, low-firing states (Klinshov et al., 2014). The rare and strong synaptic connections participate to a disproportionate degree in information processing (Nigam et al., 2016), such as feature preference and selectivity in visual cortex (Cossell et al., 2015). They may also contribute to memory recall in associative memory networks (Hiratani et al., 2013). Network simulations also indicated that lognormal-like synaptic distributions are important in the context of criticality since they support continuous transitions to chaos associated with the generation of scale-free avalanches (Kuśmierz et al., 2020). In addition, a recent computational study showed that strong synaptic inputs from the heavy tail of the lognormal synaptic efficacy distribution play a crucial role in triggering local dendritic spikes (Goetz et al., 2021) which are known to enhance nonlinear single cell computations.

## Conclusion

In sum, our work highlights the importance of a skewed, lognormal-like distribution of brain parameters. It persists through high frequency stimulation and plasticity processes and emerges even when presynaptic transmitter release is blocked. Given its importance and widespread presence in the brain, computational plasticity models should strive to maintain a skewed, lognormal-like distribution of spine sizes and synaptic weights.

## Acknowledgments

This research was supported by the German Research Foundation (Deutsche Forschungsgemeinschaft), grant number 467764793, JE 528/10-1 (to PJ), CRC1080 (to T.D.), and grant number SCHW 534/6-1 (to SWS), by the Kogge-Stiftung (to PJ) and by funds from the von Behring Röntgen Foundation (to PJ).

## Author Contributions

Conceptualization, H.C, S.W.S. and P.J.; Methodology, N.R., T.J., A.B., M.M., H.C. and P.J.; Software, N.R., T.J. and A.B.; Formal Analysis, N.R.; Investigation, N.R., A.B.; Resources, T.J., A.S., J.S.R. and N.B.; Writing – Original Draft, N.R. and P.J.; Writing – Review & Editing, N.R., T.J., A.S., A.B., M.M., J.S.R., T.D., H.C., N.B., S.W.S. and P.J; Supervision, P.J.; Project Administration, P.J.; Funding Acquisition, T.D., S.W.S. and P.J. All authors gave final approval for publication and agreed to be held accountable for the work performed therein

## Compliance with Ethical Standards

**Conflict of Interest:** The authors declare that they have no conflict of interest.

**Ethical approval: N.A.**

## Methods

### Spine data from dentate adult-born granule cells (abGCs) in rats with induced homo- and heterosynaptic plasticity

We analysed the distribution of spines in granule cell (GC) data in the dentate gyrus (DG), from Jungenitz et al. (2018). In this data set, structural homo- and heterosynaptic plasticity of spines was induced in abGCs using two hour high-frequency stimulation (HFS) of the medial perforant path (MPP) in anesthetised rats. AbGCs were stimulated at different time points after the injection of retroviral vectors (days post injection, or dpi). The cell ages used in the analysis were 21, 28 and 35 dpi. The HFS induced LTP associated with spine expansion in the middle molecular layer (MML) of the dentate gyrus (Jungenitz et al., 2018). Concurrently it induced heterosynaptic LTD associated with spine shrinkage in the inner and outer molecular layer (IML/OML).

The data set comprised spine data for individual cells in (i) the three different layers (IML, MML and OML), (ii) at the three different cell ages (21, 28 and 35 dpi), and (iii) from both the contra- and ipsilateral hemisphere. The contralateral side without the induction of synaptic plasticity (Jungenitz et al., 2018) was included as a control. All analysed spines were mushroom spines (spines with a large head in relation to the neck (Bosch and Hayashi, 2012; Rochefort and Konnerth, 2012)). Analysis was done at the level of individual cells or dentate molecular layers, separately for each layer, hemisphere, and cell age.

### Spine data from CA1 pyramidal cells in Munc13 double-knockouts

The blocked presynaptic activity data set contained spine data from CA1 pyramidal cells (PC) in hippocampal organotypic slices from Munc13 double-knockout (DKO) mice (Sigler et al., 2017). In these DKOs, the elimination of synaptic protein Munc13 causes a complete loss of spontaneous and evoked transmitter release (Varoqueaux et al., 2002). The data set comprised spine data from M13-DKOs and their controls, from three different time points of measurement (7, 14 and 21 days *in vitro*, div). The data set was split into apical and basal dendrites, and in three spine subgroups (mushroom, stubby and thin).

### Fitting a lognormal distribution to the data

The spine head area was used to analyse the distribution of spine sizes. All analyses were done with Matlab software using a custom-written script. We analysed cells individually as well as collectively by combining and averaging all cells for one condition.

From the raw data, the mean (µ) and standard deviation (σ) of the spine sizes’ natural logarithms were calculated. They functioned as a starting point for the algorithm implemented to fit the lognormal distribution over the spine data. Because the data spans multiple scales, the raw size data was normalised. For the normalisation, the integral of the spine size distribution was calculated, and the absolute number of spines in each size bin was divided by that integral.

The next step in the analysis was to build the lognormal function that would be fitted to the normalised data. For this, a customised fitting procedure had to be derived for which the probability density function (PDF) of the lognormal distribution was used:

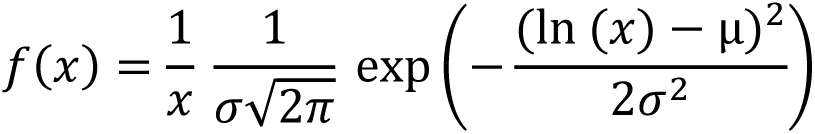

µ and σ are defined as parameters, f(x) as the dependent and x as the independent variable. The lognormal distribution was then fitted to the normalised data. With the fit function, plots and respective goodness of fit statistics for each of the fits were generated. The goodness-of-fit statistics give an indication of how well the respective fit or model fitted the data. The r-square (r^2^) value was used in all further analysis.

The key characteristic of a lognormal distribution is that the logarithm of the random variable will be normally distributed. Thus, taking the logarithm of the spine data is another good method to check if the data is distributed lognormally-like. A similar fitting procedure as above was applied. The data was first transformed by taking the logarithm of the spine sizes, then a Gaussian distribution was fitted to the logarithmic data:

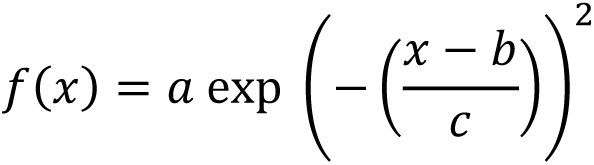

where a, b and c are the parameters, f(x) the dependent and x the independent variable. With the fit function fitting a Gaussian distribution to the logarithm of the data, new plots were generated that compared the logarithmic data with the fit.

To determine differences between the different layers, cell ages or experimental and control groups, the given r^2^ for each condition was compared, using statistical non-parametric tests. r^2^, or the coefficient of determination, is used to determine how well the variation in f(x) (the dependent variable) can be explained by x (the independent variable(s)). Essentially, it provides a measure of how well the observed outcomes can be replicated by a model. In our case, how well the applied fits describe the spine size data. The value is less than or equal to 1, with 1 being a perfect fit of the model. The coefficient of determination is calculated as follows:

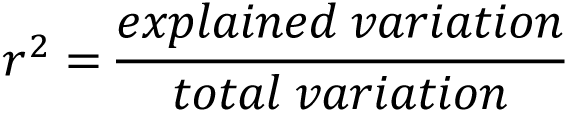

To support the findings of the goodness of fit comparisons, we also looked at the skewness (asymmetry around the mean) of the data and the width of the distribution (standard deviation of the data’s natural logarithm, in the following called sigma). More information about these comparisons can be found in the Supplementary Methods. Additionally, we conducted a model fit comparison for which we fit two additional skewed distributions (gamma and Weibull) to the data, and then used the Akaike Information Criterion (AIC) to compare all three distribution fits. This was done to see whether or not the lognormal distribution was the best fit for the data. More information about the AIC calculations and comparisons can be found in the Supplementary Methods.

### Statistical Analysis

Several statistical tests were applied to test for statistical differences of r^2^ for a lognormal and the skewness between the different conditions in both data sets. The distribution analysis showed a lognormal distribution in the spine data, so only non-parametric tests were applied.

For the hemisphere (ipsilateral / stimulated vs contralateral / non-stimulated) comparison in the rat dentate abGC spine data and the group comparison (Munc13 DKO group with blocked presynaptic release vs. control group) in the mouse CA1 PC spine data, we used a Mann-Whitney-U or ranksum test. To compare between the three different dentate layers (IML / MML / OML), we used Friedman’s test. Since all three layer-samples in one cell come from the same cell, it was a repeated measurement of multiple variables. The Kruskal-Wallis test was used for the comparison between different cell ages or cell culture ages. If significant differences (p < 0.05) were found in one sample, both for the time comparison and the layer comparison, *post-hoc* paired ranksum tests were conducted. A Bonferroni-Holm correction for multiple tests was applied to test for specific significant differences in the sample.

### Multiplicative STDP model to investigate lognormal distributions

To further investigate the influence of plasticity on the lognormal-like distribution of synaptic weights, we developed a simple model based on van Rossum et al. (2000). The model includes heterosynaptic scaling, an intrinsic multiplicative (Kesten) noise process, and an STDP learning rule with additive potentiation and multiplicative depression. The times between a presynaptic event and a postsynaptic event are written as Δt. Negative values of Δt, where the presynaptic event precedes the postsynaptic event, lead to potentiation w → w_p_ and positive values lead to depression w → w_d_.

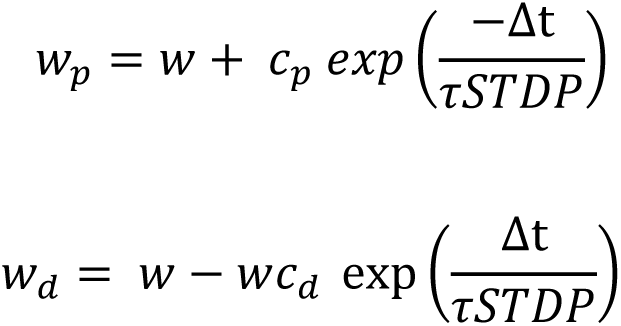

*w* is the synaptic weight, 𝑐_𝑝_ is the weight of potentiation (𝑐_𝑝_= 0.007 pS), 𝑐_𝑑_ is the weight of depression (𝑐_𝑑_ = 0.003) and 𝜏 is the time constant for STDP (𝜏𝑆𝑇𝐷𝑃 = 0.5 ms). In addition, the synapses are affected by a continuous-time multiplicative noise process of strength 4% per second. The postsynaptic neurons are modelled as leaky integrate-and-fire cells receiving 100 inputs each and uniform heterosynaptic scaling maintains a constant total conductance. The membrane time constant is 10 ms and the firing threshold is 10 mV above rest.

The model consists of a population of 1000 neurons, and the synapses are stimulated either in a Poisson manner or with periodic spiking, at different input frequencies depending on the simulation condition.

To see whether or not multiplicative noise (i.e. intrinsic mechanisms) are enough to generate a lognormal distribution, a uniform distribution was fed into the model as an initial distribution and the synaptic weights were measured throughout the simulation. The model was then used to replicate the two experimental data sets. First, the high frequency stimulation that induced LTP in the stimulated spines was recreated with the model, using periodic spiking as input at a 200 Hz frequency. This was compared with a control simulation, that received 10 Hz input. The second simulation compared intrinsic mechanisms, simulated with only multiplicative noise in a silent period, and extrinsic and intrinsic processes using a control simulation at 5 Hz receiving Poisson input. A lognormal distribution was fitted to the synaptic weight data in the same way as previously described. Additionally, the logarithm was taken of the data and a Gaussian distribution was fitted to the transformed data.

## References

Asrican, B., Lisman, J., & Otmakhov, N. (2007). Synaptic strength of individual spines correlates with bound Ca2+ –calmodulin-dependent kinase II. Journal of Neuroscience, 27(51), 14007–14011. https://doi.org/10.1523/JNEUROSCI.3587-07.2007

Barbour, B., Brunel, N., Hakim, V., & Nadal, J. P. (2007). What can we learn from synaptic weight distributions? Trends in Neurosciences, 30(12), 622–629. https://doi.org/10.1016/j.tins.2007.09.005

Beining, M., Jungenitz, T., Radic, T., Deller, T., Cuntz, H., Jedlicka, P., & Schwarzacher, S. W. (2017). Adult-born dentate granule cells show a critical period of dendritic reorganization and are distinct from developmentally born cells. Brain structure & function, 222(3), 1427–1446. https://doi.org/10.1007/s00429-016-1285-y

Bhatt, D. H., Zhang, S., & Gan, W. B. (2009). Dendritic spine dynamics. Annual Review of Physiology, 71, 261–282. https://doi.org/10.1146/annurev.physiol.010908.163140

Bosch, M., & Hayashi, Y. (2012). Structural plasticity of dendritic spines. Current opinion in neurobiology, 22(3), 383–388. https://doi.org/10.1016/j.conb.2011.09.002

Bourne, J., & Harris, K. M. (2007). Do thin spines learn to be mushroom spines that remember? Current Opinion in Neurobiology, 17(3), 381–386. https://doi.org/10.1016/j.conb.2007.04.009

Bromer, C., Bartol, T. M., Bowden, J. B., Hubbard, D. D., Hanka, D. C., Gonzalez, P. v., Kuwajima, M., Mendenhall, J. M., Parker, P. H., Abraham, W. C., Sejnowski, T. J., & Harris, K. M. (2018). Long-term potentiation expands information content of hippocampal dentate gyrus synapses. Proceedings of the National Academy of Sciences of the United States of America, 115(10), E2410–E2418. https://doi.org/10.1073/pnas.1716189115

Buzsáki, G., & Mizuseki, K. (2014). The log-dynamic brain: How skewed distributions affect network operations. Nature Reviews Neuroscience, 15(4), 264–278. Nature Publishing Group. https://doi.org/10.1038/nrn3687

Cossell, L., Iacaruso, M. F., Muir, D. R., Houlton, R., Sader, E. N., Ko, H., Hofer, S. B., & Mrsic-Flogel, T. D. (2015). Functional organization of excitatory synaptic strength in primary visual cortex. Nature, 518(7539), 399–403. https://doi.org/10.1038/nature14182

de Vivo, L., Bellesi, M., Marshall, W., Bushong, E. A., Ellisman, M. H., Tononi, G., & Cirelli, C. (2017). Ultrastructural evidence for synaptic scaling across the wake/sleep cycle. Science, 355(6324), 507–510. https://doi.org/10.1126/science.aah5982

Dorkenwald, S., Turner, N. L., Macrina, T., Lee, K., Lu, R., Wu, J., Bodor, A. L., Bleckert, A. A., Brittain, D., Kemnitz, N., Silversmith, W. M., Ih, D., Zung, J., Zlateski, A., Tartavull, I., Yu, S. C., Popovych, S., Wong, W., Castro, M. et al. (2019). Binary and analog variation of synapses between cortical pyramidal neurons. BioRxiv. https://doi.org/10.1101/2019.12.29.890319

Effenberger, F., Jost, J., & Levina, A. (2015). Self-organization in balanced state networks by STDP and homeostatic plasticity. PLOS Computational Biology, 11(9), e1004420. https://doi.org/10.1371/journal.pcbi.1004420

Gal, E., London, M., Globerson, A., Ramaswamy, S., Reimann, M. W., Muller, E., Markram, H., & Segev, I. (2017). Rich cell-type-specific network topology in neocortical microcircuitry. Nature Neuroscience, 20(7), 1004–1013. https://doi.org/10.1038/nn.4576

Gilson, M., & Fukai, T. (2011). Stability versus neuronal specialization for STDP: long-tail weight distributions solve the dilemma. PLoS ONE, 6(10), e25339. https://doi.org/10.1371/journal.pone.0025339

Goetz, L., Roth, A., & Häusser, M. (2021). Active dendrites enable strong but sparse inputs to determine orientation selectivity. Proceedings of the National Academy of Sciences, 118(30), e2017339118. https://doi.org/10.1073/pnas.2017339118

Harms, K. J., & Craig, A. M. (2005). Synapse composition and organization following chronic activity blockade in cultured hippocampal neurons. Journal of Comparative Neurology, 490(1), 72–84. https://doi.org/10.1002/cne.20635

Harms, K. J., Tovar, K. R., & Craig, A. M. (2005). Synapse-specific regulation of AMPA receptor subunit composition by activity. Journal of Neuroscience, 25(27), 6379– 6388. https://doi.org/10.1523/JNEUROSCI.0302-05.2005

Harris, K. M. (2020). Structural LTP: from synaptogenesis to regulated synapse enlargement and clustering. Current Opinion in Neurobiology, 63, 189–197. https://doi.org/10.1016/j.conb.2020.04.009

Harris, K. M., Jensen, F. E., & Tsao, B. (1992). Three-dimensional structure of dendritic spines and synapses in rat hippocampus (CA1) at postnatal day 15 and adult ages: Implications for the maturation of synaptic physiology and long-term potentiation. Journal of Neuroscience, 12(8), 2685–2705.

Harris, K. M., & Stevens, J. K. (1989). Dendritic spines of CA 1 pyramidal cells in the rat hippocampus: serial electron microscopy with reference to their biophysical characteristics. The Journal of Neuroscience, 9(8), 2982–2997. https://doi.org/10.1523/JNEUROSCI.09-08-02982.1989

Hazan, L., & Ziv, N. E. (2020). Activity dependent and independent determinants of synaptic size diversity. Journal of Neuroscience, 40(14), 2828–2848. https://doi.org/10.1523/JNEUROSCI.2181-19.2020

Held, R.G., Liu, C., Ma, K., Ramsey, A.M., Tarr, T.B., De Nola, G., Wang, S.S.H., Wang, J., van den Maagdenberg, A.M.J.M., Schneider, T., Sun, J., Blanpied, T.A., Kaeser, P.S. (2020). Synapse and Active Zone Assembly in the Absence of Presynaptic Ca2+ Channels and Ca2+ Entry. Neuron, 107(4), 667–683.e9. https://doi.org/10.1016/j.neuron.2020.05.032

Hiratani, N., Teramae, J. N., & Fukai, T. (2013). Associative memory model with long-tail-distributed Hebbian synaptic connections. Frontiers in Computational Neuroscience, 6, 102. https://doi.org/10.3389/fncom.2012.00102

Hung, A. Y., Futai, K., Sala, C., Valtschanoff, J. G., Ryu, J., Woodworth, M. A., Kidd, F. L., Sung, C. C., Miyakawa, T., Bear, M. F., Weinberg, R. J., & Sheng, M. (2008). Smaller dendritic spines, weaker synaptic transmission, but enhanced spatial learning in mice lacking Shank1. Journal of Neuroscience, 28(7), 1697–1708. https://doi.org/10.1523/JNEUROSCI.3032-07.2008

Ikegaya, Y., Sasaki, T., Ishikawa, D., Honma, N., Tao, K., Takahashi, N., Minamisawa, G., Ujita, S., & Matsuki, N. (2013). Interpyramid spike transmission stabilizes the sparseness of recurrent network activity. Cerebral Cortex, 23(2), 293–304. https://doi.org/10.1093/cercor/bhs006

Iyer, R., Menon, V., Buice, M., Koch, C., & Mihalas, S. (2013). The influence of synaptic weight distribution on neuronal population dynamics. PLoS Computational Biology, 9(10), e1003248. https://doi.org/10.1371/journal.pcbi.1003248

Jungenitz, T., Beining, M., Radic, T., Deller, T., Cuntz, H., Jedlicka, P., & Schwarzacher, S. W. (2018). Structural homo- and heterosynaptic plasticity in mature and adult newborn rat hippocampal granule cells. Proceedings of the National Academy of Sciences of the United States of America, 115(20), E4670–E4679. https://doi.org/10.1073/pnas.1801889115

Kasai, H., Fukuda, M., Watanabe, S., Hayashi-Takagi, A., & Noguchi, J. (2010). Structural dynamics of dendritic spines in memory and cognition. Trends in Neurosciences, 33(3), 121–129. https://doi.org/10.1016/j.tins.2010.01.001

Kasai, H., Matsuzaki, M., Noguchi, J., Yasumatsu, N., & Nakahara, H. (2003). Structure-stability-function relationships of dendritic spines. Trends in Neurosciences, 26(7), 360–368. https://doi.org/10.1016/S0166-2236(03)00162-0

Kasai, H., Ziv, N. E., Okazaki, H., Yagishita, S., & Toyoizumi, T. (2021). Spine dynamics in the brain, mental disorders and artificial neural networks. Nature Reviews Neuroscience, 22(7), 407–422. https://doi.org/10.1038/s41583-021-00467-3

Kleinjan, M. S., Buchta, W. C., Ogelman, R., Hwang, I. W., Kuwajima, M., Hubbard, D. D., Kareemo, D. J., Prikhodko, O., Olah, S. L., Gomez Wulschner, L. E., Abraham, W. C., Franco, S. J., Harris, K. M., Oh, W. C., & Kennedy, M. J. (2023). Dually innervated dendritic spines develop in the absence of excitatory activity and resist plasticity through tonic inhibitory crosstalk. Neuron, 111(3), 362–371.e6. https://doi.org/10.1016/j.neuron.2022.11.002

Klinshov, V. v., Teramae, J., Nekorkin, V. I., & Fukai, T. (2014). Dense neuron clustering explains connectivity statistics in cortical microcircuits. PLoS ONE, 9(4), e94292. https://doi.org/10.1371/journal.pone.0094292

Kuśmierz, Ł., Ogawa, S., & Toyoizumi, T. (2020). Edge of chaos and avalanches in neural networks with heavy-tailed synaptic weight distribution. Physical Review Letters, 125(2), 028101. https://doi.org/10.1103/PhysRevLett.125.028101

Lefort, S., Tomm, C., Floyd Sarria, J.-C., & Petersen, C. C. H. (2009). The excitatory neuronal network of the C2 barrel column in mouse primary somatosensory cortex. Neuron, 61(2), 301–316. https://doi.org/10.1016/j.neuron.2008.12.020

Loewenstein, Y., Kuras, A., & Rumpel, S. (2011). Multiplicative dynamics underlie the emergence of the log-normal distribution of spine sizes in the neocortex in vivo. Journal of Neuroscience, 31(26), 9481–9488. https://doi.org/10.1523/JNEUROSCI.6130-10.2011

Lu, W., Bushong, E. A., Shih, T. P., Ellisman, M. H., & Nicoll, R. A. (2013). The cell-autonomous role of excitatory synaptic transmission in the regulation of neuronal structure and function. Neuron, 78(3), 433–439. https://doi.org/10.1016/j.neuron.2013.02.030

Mateos, J. M., Lüthi, A., Savic, N., Stierli, B., Streit, P., Gähwiler, B. H., & McKinney, R. A. (2007). Synaptic modifications at the CA3-CA1 synapse after chronic AMPA receptor blockade in rat hippocampal slices. The Journal of Physiology, 581(1), 129–138. https://doi.org/10.1113/jphysiol.2006.120550

Matsuzaki, M., Ellis-Davies, G., Nemoto, T., Miyashita, Y., Iino, M., & Kasai, H. (2001). Dendritic spine geometry is critical for AMPA receptor expression in hippocampal CA1 pyramidal neurons. Nature Neuroscience, 4, 1086–1092. https://doi.org/10.1038/nn736

McKinney, R. A. (2010). Excitatory amino acid involvement in dendritic spine formation, maintenance and remodelling. Journal of Physiology, 588(1), 107–116. https://doi.org/10.1113/jphysiol.2009.178905

McKinney, R. A., Capogna, M., Dürr, R., Gähwiler, B. H., & Thompson, S. M. (1999). Miniature synaptic events maintain dendritic spines via AMPA receptor activation. Nature Neuroscience, 2(1), 44–49. https://doi.org/10.1038/4548

Merchán-Pérez, A., Rodríguez, J. R., González, S., Robles, V., Defelipe, J., Larrañaga, P., & Bielza, C. (2014). Three-dimensional spatial distribution of synapses in the neocortex: A dual-beam electron microscopy study. Cerebral Cortex, 24(6), 1579– 1588. https://doi.org/10.1093/cercor/bht018

Minerbi, A., Kahana, R., Goldfeld, L., Kaufman, M., Marom, S., & Ziv, N. E. (2009). Long-term relationships between synaptic tenacity, synaptic remodeling, and network activity. PLoS Biology, 7(6), e1000136. https://doi.org/10.1371/journal.pbio.1000136

Mizuseki, K., & Buzsáki, G. (2013). Preconfigured, skewed distribution of firing rates in the hippocampus and entorhinal cortex. Cell Reports, 4(5), 1010–1021. https://doi.org/10.1016/j.celrep.2013.07.039

Mongillo, G., Rumpel, S., & Loewenstein, Y. (2017). Intrinsic volatility of synaptic connections — a challenge to the synaptic trace theory of memory. Current Opinion in Neurobiology, 46, 7–13). https://doi.org/10.1016/j.conb.2017.06.006

Montero-Crespo, M., Dominguez-Alvaro, M., Rondon-Carrillo, P., Alonso-Nanclares, L., Defelipe, J., & Blazquez-Llorca, L. (2020). Three-dimensional synaptic organization of the human hippocampal ca1 field. ELife, 9, e57013. https://doi.org/10.7554/eLife.57013

Nigam, S., Shimono, M., Ito, S., Yeh, F. C., Timme, N., Myroshnychenko, M., Lapish, C. C., Tosi, Z., Hottowy, P., Smith, W. C., Masmanidis, S. C., Litke, A. M., Sporns, O., & Beggs, J. M. (2016). Rich-club organization in effective connectivity among cortical neurons. Journal of Neuroscience, 36(3), 655–669. https://doi.org/10.1523/JNEUROSCI.2177-15.2016

Nishiyama, J., & Yasuda, R. (2015). Biochemical computation for spine structural plasticity. Neuron, 87(1), 63–75. https://doi.org/10.1016/j.neuron.2015.05.043

Nusser, Z., Lujan, R., Laube, G., Roberts, J. D. B., Molnar, E., & Somogyi, P. (1998). Cell type and pathway dependence of synaptic AMPA receptor number and variability in the hippocampus. Neuron, 21(3), 545–559. https://doi.org/10.1016/S0896-6273(00)80565-6

Ohkawa, N., Saitoh, Y., Tokunaga, E., Nihonmatsu, I., Ozawa, F., Murayama, A., Shibata, F., Kitamura, T., & Inokuchi, K. (2012). Spine formation pattern of adult-born neurons is differentially modulated by the induction timing and location of hippocampal plasticity. PloS one, 7(9), e45270. https://doi.org/10.1371/journal.pone.0045270

Omura, Y., Carvalho, M. M., Inokuchi, K., & Fukai, T. (2015). A lognormal recurrent network model for burst generation during hippocampal sharp waves. Journal of Neuroscience, 35(43), 14585–14601. https://doi.org/10.1523/JNEUROSCI.4944-14.2015

Radic, T., Jungenitz, T., Singer, M., Beining, M., Cuntz, H., Vlachos, A., Deller, T., & Schwarzacher, S. W. (2017). Time-lapse imaging reveals highly dynamic structural maturation of postnatally born dentate granule cells in organotypic entorhino-hippocampal slice cultures. Scientific Reports, 7, 43724. https://doi.org/10.1038/srep43724

Rochefort, N. L., & Konnerth, A. (2012). Dendritic spines: from structure to in vivo function. EMBO reports, 13(8), 699–708. https://doi.org/10.1038/embor.2012.102

Sando, R., Bushong, E., Zhu, Y., Huang, M., Considine, C., Phan, S., Ju, S., Uytiepo, M., Ellisman, M., & Maximov, A. (2017). Assembly of excitatory synapses in the absence of glutamatergic neurotransmission. Neuron, 94(2), 312–321. https://doi.org/10.1016/j.neuron.2017.03.047

Santuy, A., Rodríguez, J. R., DeFelipe, J., & Merchán-Pérez, A. (2018). Study of the size and shape of synapses in the juvenile rat somatosensory cortex with 3D electron microscopy. ENeuro, 5(1), ENEURO.0377-17.2017. https://doi.org/10.1523/ENEURO.0377-17.2017

Scheler, G. (2017). Logarithmic distributions prove that intrinsic learning is Hebbian. F1000Research, 6, 1222. https://doi.org/10.12688/f1000research.12130.1

Segal, M. (2017). Dendritic spines: Morphological building blocks of memory. Neurobiology of Learning and Memory, 138, 3–9. https://doi.org/10.1016/j.nlm.2016.06.007

Shomar, A., Geyrhofer, L., Ziv, N. E., & Brenner, N. (2017). Cooperative stochastic binding and unbinding explain synaptic size dynamics and statistics. PLoS Computational Biology, 13(7), e1005668. https://doi.org/10.1371/journal.pcbi.1005668

Sigler, A., Oh, W. C., Imig, C., Altas, B., Kawabe, H., Cooper, B. H., Kwon, H. B., Rhee, J. S., & Brose, N. (2017). Formation and maintenance of functional spines in the absence of presynaptic glutamate release. Neuron, 94(2), 304–311.e4. https://doi.org/10.1016/j.neuron.2017.03.029

Song, S., Sjöström, P. J., Reigl, M., Nelson, S., & Chklovskii, D. B. (2005). Highly nonrandom features of synaptic connectivity in local cortical circuits. PLoS Biology, 3(3), e68. https://doi.org/10.1371/journal.pbio.0030068

Sorra, K. E., & Harris, K. M. (1998). Stability in synapse number and size at 2 Hr after long-term potentiation in hippocampal area CA1. Journal of Neuroscience, 18(2), 658–671. https://doi.org/10.1523/jneurosci.18-02-00658.1998

Statman, A., Kaufman, M., Minerbi, A., Ziv, N. E., & Brenner, N. (2014). Synaptic size dynamics as an effectively stochastic process. PLoS Computational Biology, 10(10), e1003846. https://doi.org/10.1371/journal.pcbi.1003846

Suratkal, S. S., Yen, Y. H., & Nishiyama, J. (2021). Imaging dendritic spines: molecular organization and signaling for plasticity. Current Opinion in Neurobiology, 67, 66–74. https://doi.org/10.1016/j.conb.2020.08.006

Takumi, Y., Ramírez-León, V., Laake, P., Rinvik, E., & Ottersen, O. P. (1999). Different modes of expression of AMPA and NMDA receptors in hippocampal synapses. Nature Neuroscience, 2(7), 618–624. https://doi.org/10.1038/10172

Teramae, J. N., & Fukai, T. (2014). Computational implications of lognormally distributed synaptic weights. Proceedings of the IEEE, 102(4), 500–512. https://doi.org/10.1109/JPROC.2014.2306254

Teramae, J. N., Tsubo, Y., & Fukai, T. (2012). Optimal spike-based communication in excitable networks with strong-sparse and weak-dense links. Scientific Reports, 2, 485. https://doi.org/10.1038/srep00485

Triesch, J., Vo, A. D., & Hafner, A. S. (2018). Competition for synaptic building blocks shapes synaptic plasticity. ELife, 7, e37836. https://doi.org/10.7554/eLife.37836

van Huizen, F., Romijn, H. J., & Habets, A. M. (1985). Synaptogenesis in rat cerebral cortex cultures is affected during chronic blockade of spontaneous bioelectric activity by tetrodotoxin. Developmental Brain Research, 351(1), 67–80. https://doi.org/10.1016/0165-3806(85)90232-9

van Rossum, M. C. W., Bi, G. Q., & Turrigiano, G. G. (2000). Stable Hebbian learning from spike timing-dependent plasticity. Journal of Neuroscience, 20(23), 8812– 8821. https://doi.org/10.1523/jneurosci.20-23-08812.2000

Varoqueaux, F., Sigler, A., Rhee, J. S., Brose, N., Enk, C., Reim, K., & Rosenmund, C. (2002). Total arrest of spontaneous and evoked synaptic transmission but normal synaptogenesis in the absence of Munc13-mediated vesicle priming. Proceedings of the National Academy of Sciences of the United States of America, 99(13), 9037–9042. https://doi.org/10.1073/pnas.122623799

Yasumatsu, N., Matsuzaki, M., Miyazaki, T., Noguchi, J., & Kasai, H. (2008). Principles of long-term dynamics of dendritic spines. Journal of Neuroscience, 28(50), 13592– 13608. https://doi.org/10.1523/JNEUROSCI.0603-08.2008

Yap, K., Drakew, A., Smilovic, D., Rietsche, M., Paul, M. H., Vuksic, M., Del Turco, D., & Deller, T. (2020). The actin-modulating protein synaptopodin mediates long-term survival of dendritic spines. eLife, 9, e62944. https://doi.org/10.7554/eLife.62944

Zheng, P., Dimitrakakis, C., & Triesch, J. (2013). Network self-organization explains the statistics and dynamics of synaptic connection strengths in cortex. PLoS Computational Biology, 9(1), e1002848. https://doi.org/10.1371/journal.pcbi.1002848

Ziff, E.B., (1997) Enlightening the postsynaptic density. Neuron, 19(6), 1163 – 1174. https://doi.org/10.1016/S0896-6273(00)80409-2

Zito, K., Scheuss, V., Knott, G., Hill, T., & Svoboda, K. (2009). Rapid functional maturation of nascent dendritic spines. Neuron, 61(2), 247–258. https://doi.org/10.1016/j.neuron.2008.10.054

Ziv, N. E., & Brenner, N. (2018). Synaptic tenacity or lack thereof: Spontaneous remodeling of synapses. Trends in Neurosciences, 41(2), 89–99. https://doi.org/10.1016/j.tins.2017.12.003

