## Supplementary Material for "Skewed distribution of spines is independent of presynaptic transmitter release and synaptic plasticity and emerges early during adult neurogenesis"

*Other distributions and Akaike information criterion (AIC) comparison*

We compared two other skewed distributions with the lognormal distribution, to see how well those fit the data and to determine if the lognormal distribution is indeed the best fit to describe the data. We chose the gamma and Weibull distributions for their similarities to the lognormal distribution in shapes and parameters.

The Gamma distribution is described by two parameters: the shape parameter k and scale parameter θ (sometimes referred to as α and β respectively). The probability density function that was used for the fit was the following:

$$f(x) = \frac{1}{\Gamma(k)\theta^k} x^{k-1} e^{-\frac{x}{\theta}}$$

The Weibull distribution is described by two parameters as well: the scale parameter λ and shape parameter k. The probability density function for the Weibull distribution fit is the following:

$$f(x) = \frac{b}{a}\left(\frac{x}{a}\right)^{b-1} e^{-\left(\frac{x}{a}\right)^b}$$

In order to compare the three distributions, we used the Akaike Information Criterion (AIC). The AIC compares the quality of a given set of models, in our case a different set of probability distributions and fits, with each other. It takes the number of parameters used for the model into consideration to avoid overfitting. In our case, all three candidate models have two parameters, so the penalty for having more parameters doesn’t play a role. The AIC in this case still offers a principled way of comparing the relative generality of the fitted distributions. However, the AIC doesn’t give any information about the absolute quality of the model, only the relative quality of a model for a given data set compared to other models. The AIC is calculated as follows:

$$AIC = 2k - 2 * \ln(L)$$

k being the number of parameters and L the maximum value of the likelihood function. L was calculated using the ‘maximum likelihood estimation’ (short mle) function in MATLAB to calculate the maximum likelihood estimates for the parameters of the respective function. These estimates, together with the spine size data, were then fed into the probability density function of each distribution respectively, to calculate the y-values for the maximum likelihood function. The logarithm was taken of the y-values and summed up to get the maximum value of the likelihood function, L. With this, the AIC could be calculated for each distribution, and then compared to find the distribution that describes the spine data the best. The lower the AIC value the better the model fits the data.

Supplementary figures 5, 6 and 11 show the AIC comparisons between the three different distributions. The AIC has a high variability in cell individual comparisons, but on average the lognormal distribution has a slight advantage over the other two distributions, both in the abGCs ipsilaterally (figure S5) and contralaterally (figure S6) and the CA1 PNs mushroom spines in both experimental conditions (figure S11).

*Skewness and Sigma as additional quantification*

In addition to goodness of fit as quantification method for the lognormal fit, we used the skewness of the data. Skewness is a measure of asymmetry around the mean of a probability distribution of a random variable. A positive skew typically implies a longer tail on the right side of the distribution, with most data accumulating on the left side. This distribution then is referred to as right-skewed or right-tailed. The lognormal distribution is an example of such a distribution. If a distribution is symmetric, meaning that the mean is equal to the median, it has zero skewness, such as in a normal / Gaussian distribution (although the converse does not necessarily hold). The skewness of the raw data was calculated with the following equation:

$$s = E\left[\left(\frac{x - \mu}{\sigma}\right)^3\right]$$

μ represents the mean of the data, σ is the standard deviation of the data and E(t) is the expected value of the quantity t. This calculation was applied to the spine data of both rat abGCs as well as murine Munc13 DKO CA1 PCs to firstly exclude the Gaussian distribution as a possible fit for the data and secondly to acquire an asymmetry measure of the probability distribution.

The width of the distribution around the peak is quantified with the standard deviation of the variables' natural logarithm, in the following labelled as sigma. Sigma defines the shape of the distribution and indicates the range of overall spine sizes. Sigma was calculated for each distribution during the lognormal fitting procedure by taking the logarithm of the standard deviation. We then compared sigma values in both data sets, in the same fashion as was done with goodness of fit and skewness values.

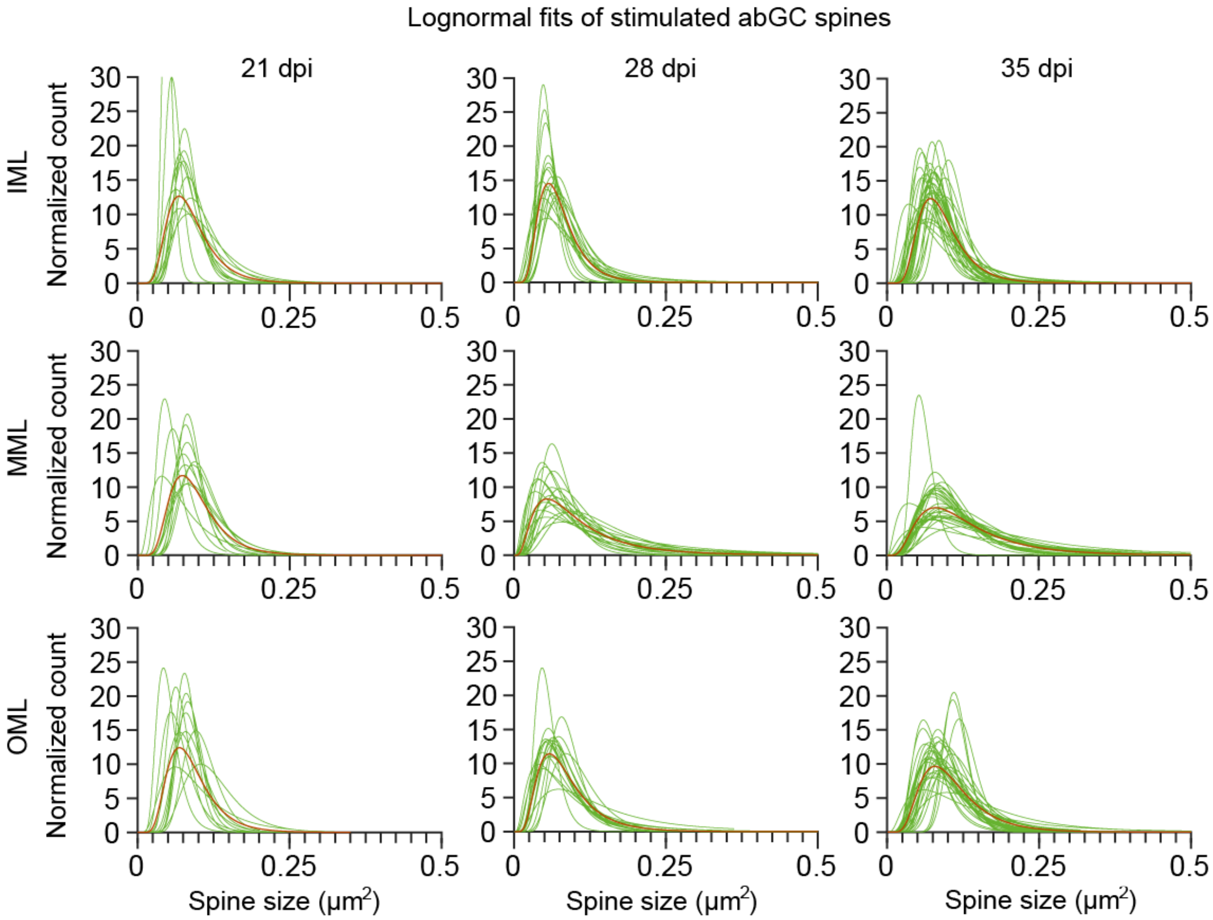


***Supplementary Figure 1. Logfits for spine size data from individual neurons (abGCs) in each layer and at each cell age in the ipsilateral (stimulated) hemisphere of anesthetised rats.*** *Each individual neuron (shown in green) shows a lognormal-like, skewed distribution. The red fit shows the average distribution that can be seen in Figure 1.*

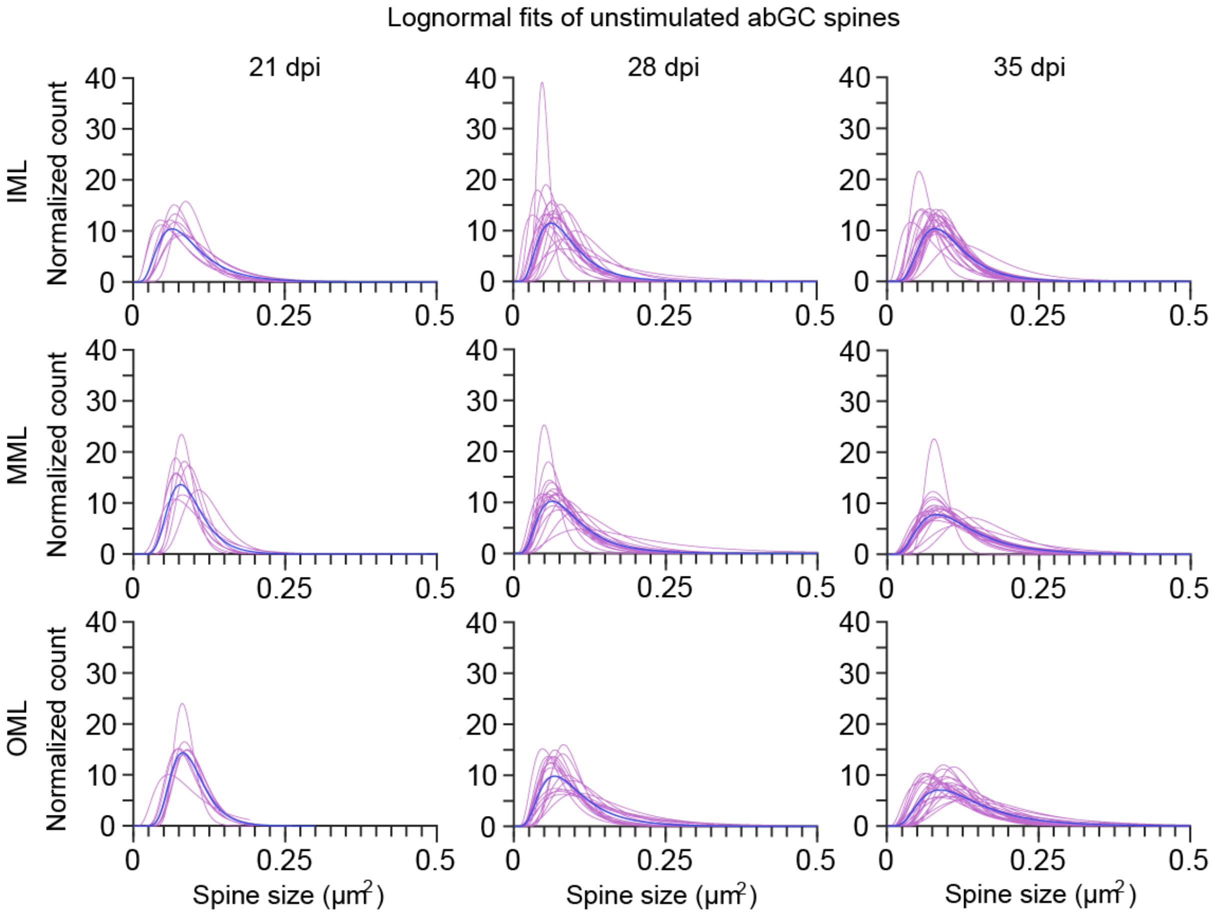


**Supplementary Figure 2.** ***Logfits for spine size data from individual neurons (abGCs) in each layer and at each time point in the contralateral hemisphere of anesthetised rats.*** *Each individual neuron (shown in magenta) shows a lognormal-like, skewed distribution. The blue fit shows the average distribution that can be seen in Figure 1.*

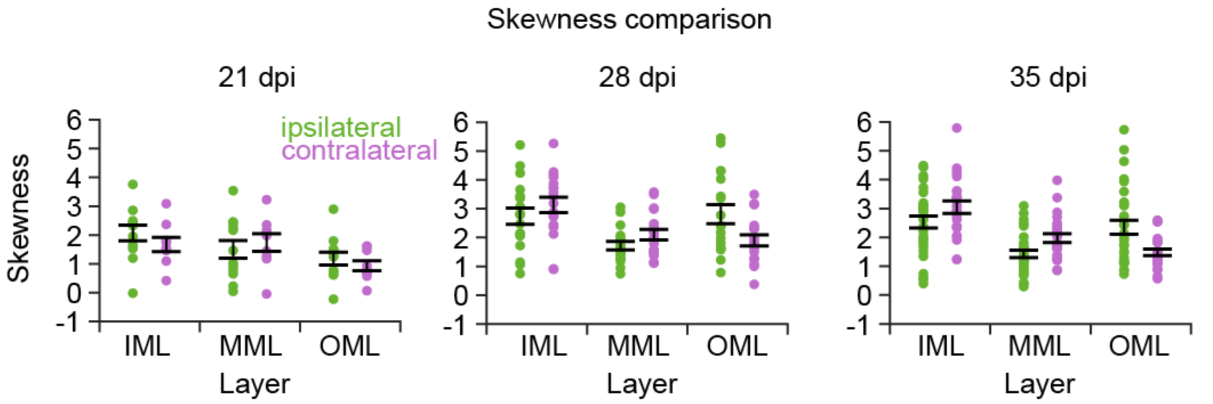


***Supplementary Figure 3. The skewness analysis reveals the spine data is positively skewed and not a symmetrical distribution.*** *The skewness values show a similar result to the goodness of fit and are similar in the ipsilateral (stimulated) and contralateral (control) dentate gyrus layers. Left, middle,*

*right panel: 21, 28 and 35 dpi, respectively. Left, middle, right panel: 21, 28 and 35 dpi, respectively. Each dot represents a single cell. The error bar represents SEM.*

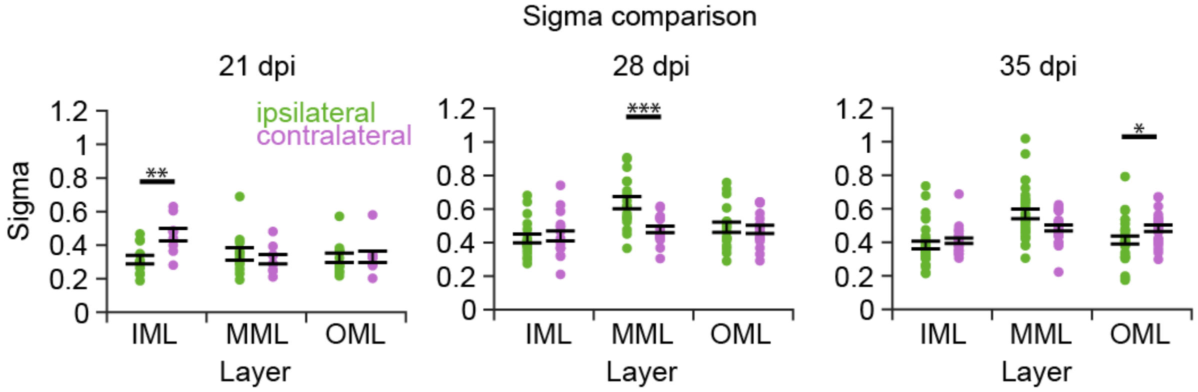


***Supplementary Figure 4. The sigma value comparison of the abGC data set reveals small changes and differences between the stimulated ipsilateral side and the unstimulated contralateral hemisphere.*** *There are significant differences ($p < 0.05$) in the IML at 21 dpi, in the MML at 28 dpi and in the OML at 35 dpi. In the case of the MML, the width of the distribution increases significantly for the ipsilateral hemisphere due to the stimulation increasing the range of spine sizes in correspondence with homosynaptic LTP (a trend for this can be seen at 35 dpi as well in the MML, however it is not significant). In the IML and OML, the opposite happens, and the contralateral side increases in width, in correspondence to heterosynaptic LTD.*

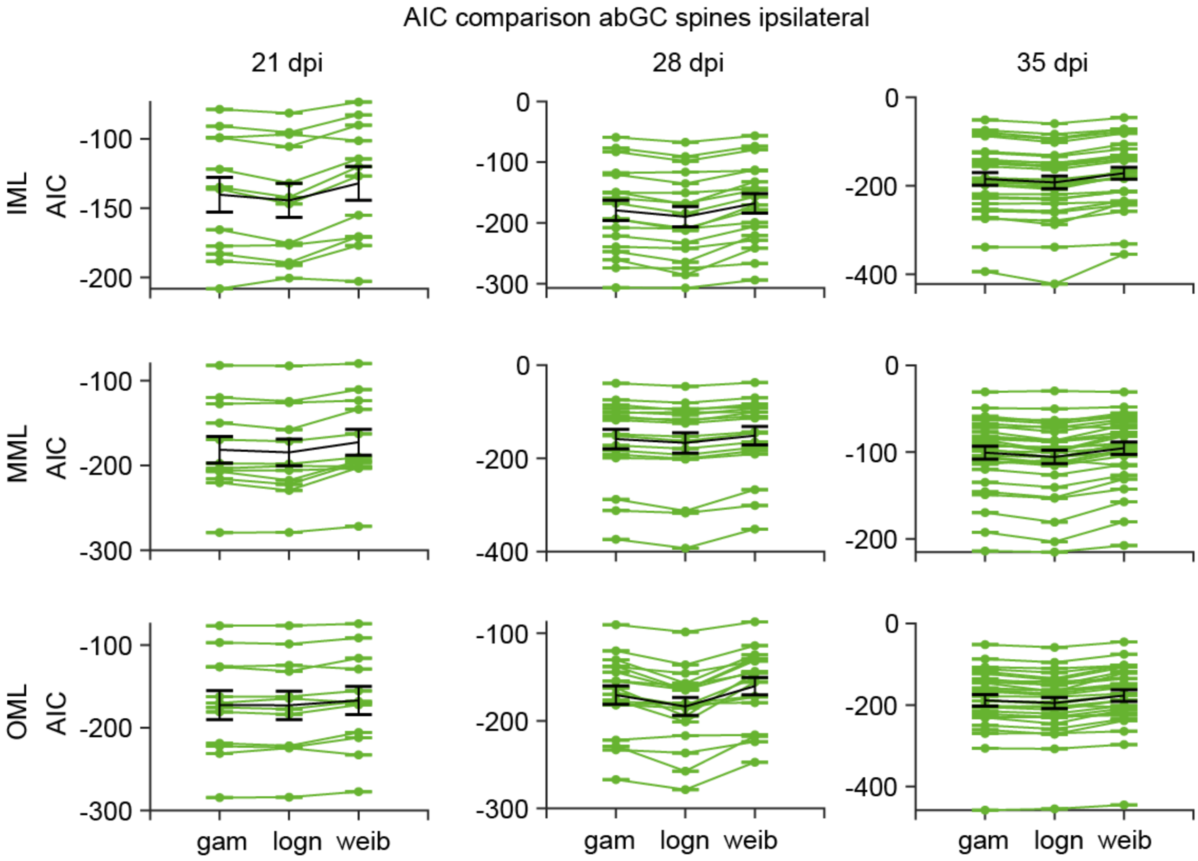


***Supplementary Figure 5. Akaike Information Criterion (AIC) in individual cells of the spine size data from anesthetised rat abGCs ipsilaterally shows a small advantage of the lognormal distribution as the best model fit for the data compared to other skewed distributions.*** *In all layers and at all cell ages, the lognormal distribution shows a slightly lower AIC value compare to other distributions. The lower the AIC, the better the fit of the model. In the cell specific analyses, in most comparisons, the lognormal distribution has the best fit. Each line represents one cell. gam – gamma distribution, logn – lognormal distribution, weib – Weibull distribution*

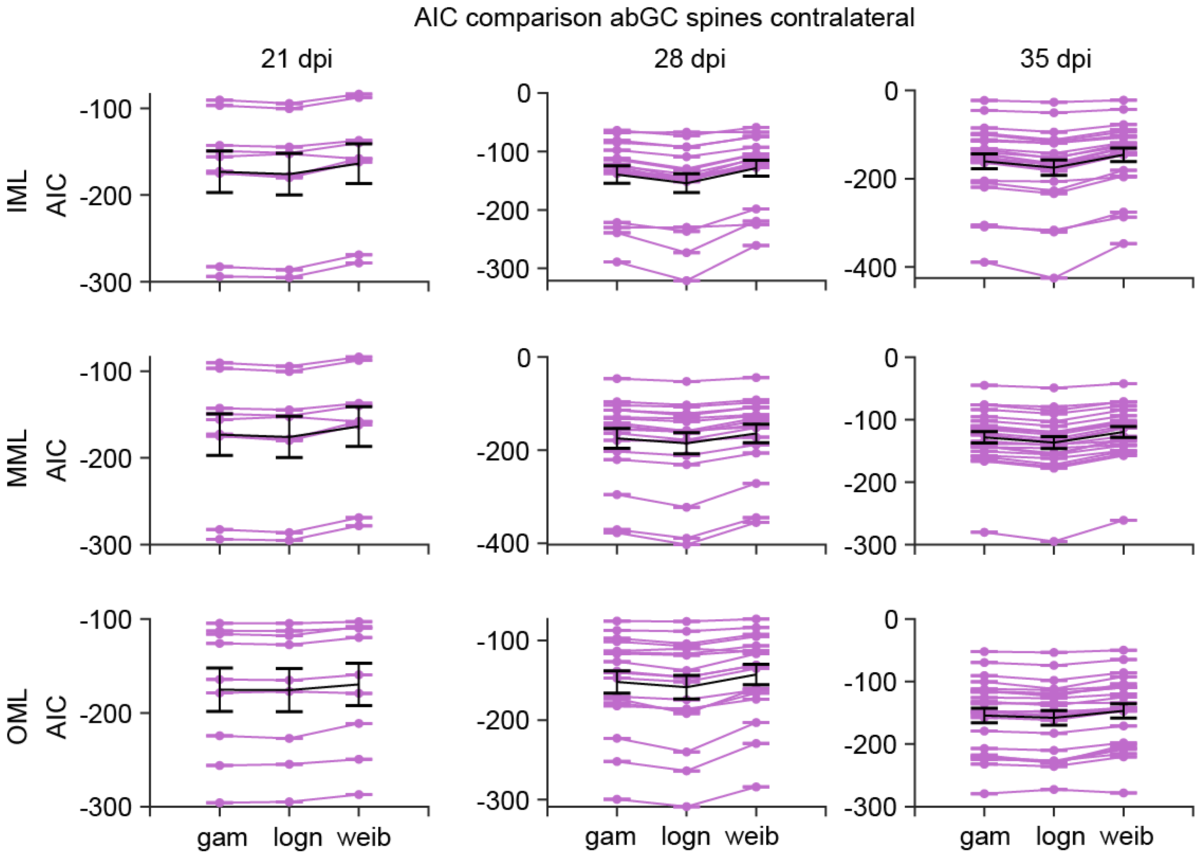


***Supplementary Figure 6. Akaike Information Criterion (AIC) in individual cells of the spine size data from anesthetised rat abGCs contralaterally shows a small advantage of the lognormal distribution as the best model fit for the data compared to other skewed distributions.*** *In all layers and at all cell ages, the lognormal distribution shows a slightly lower AIC value compare to other distributions. The lower the AIC, the better the fit of the model. In the cell specific analyses, in most comparisons, the lognormal distribution has the best fit, especially at 28 and 35 dpi. Each line represents one cell. gam – gamma distribution, logn – lognormal distribution, weib – Weibull distribution*

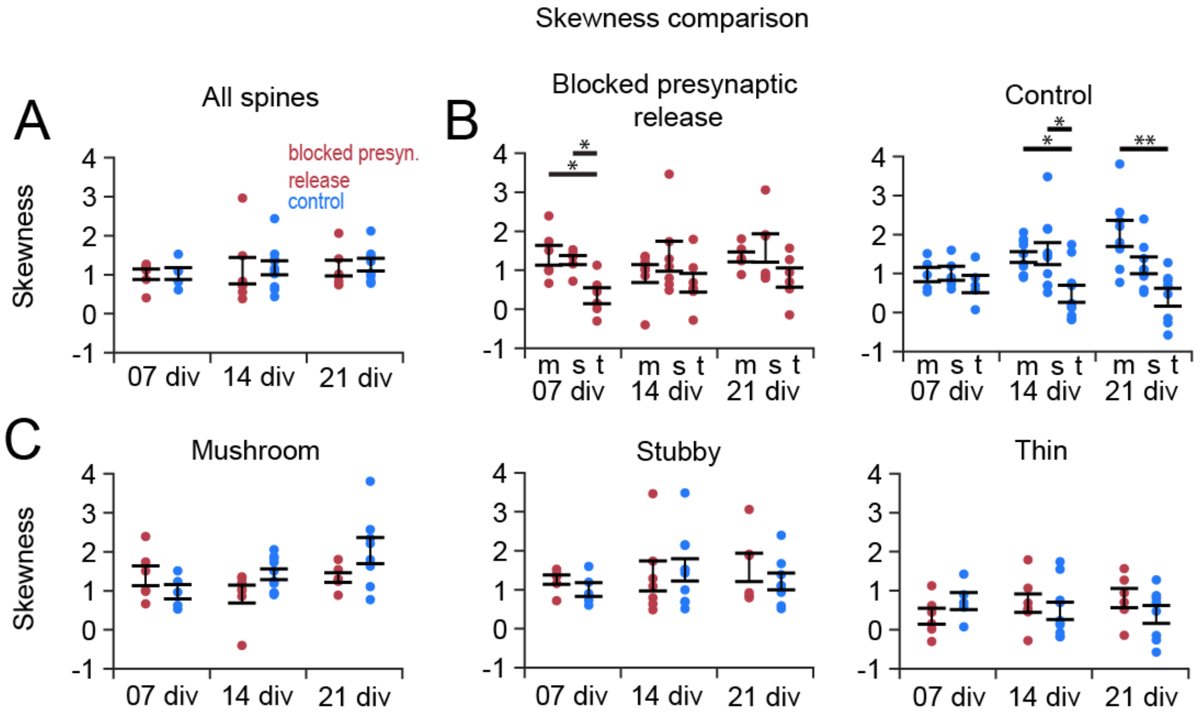


***Supplementary Figure 7. The skewness comparison in individual CA1 PCs from Munc-13 DKO (blocked presynaptic release) and WT (control) organotypic slice cultures, for all spine subtypes together and separately.*** *(A) The skewness values for all spine types were similar in the blocked presynaptic group and the control group. There are no significant differences between the groups or between the time points. Both groups show a positive skew at all cell ages. (B) Comparing the skewness of the different spine subgroups, in the blocked presynaptic release group (left panel) and the control group (right panel). Thin spines show the smallest values, with some cells even showing values below zero, indicating a skewness to the left. (C) Analysis and comparison of the skewness for each spine subtype separately. All mushroom (left panel) and stubby (middle panel) spines show a positive skewness with one cell in the mushroom comparison that has a negative skew. Thin spines show a lower skewness in general, with multiple cells having zero or lower skewness, indicating either a symmetric or skewness to the left. There are no significant differences between the two experimental groups in either spine type.*

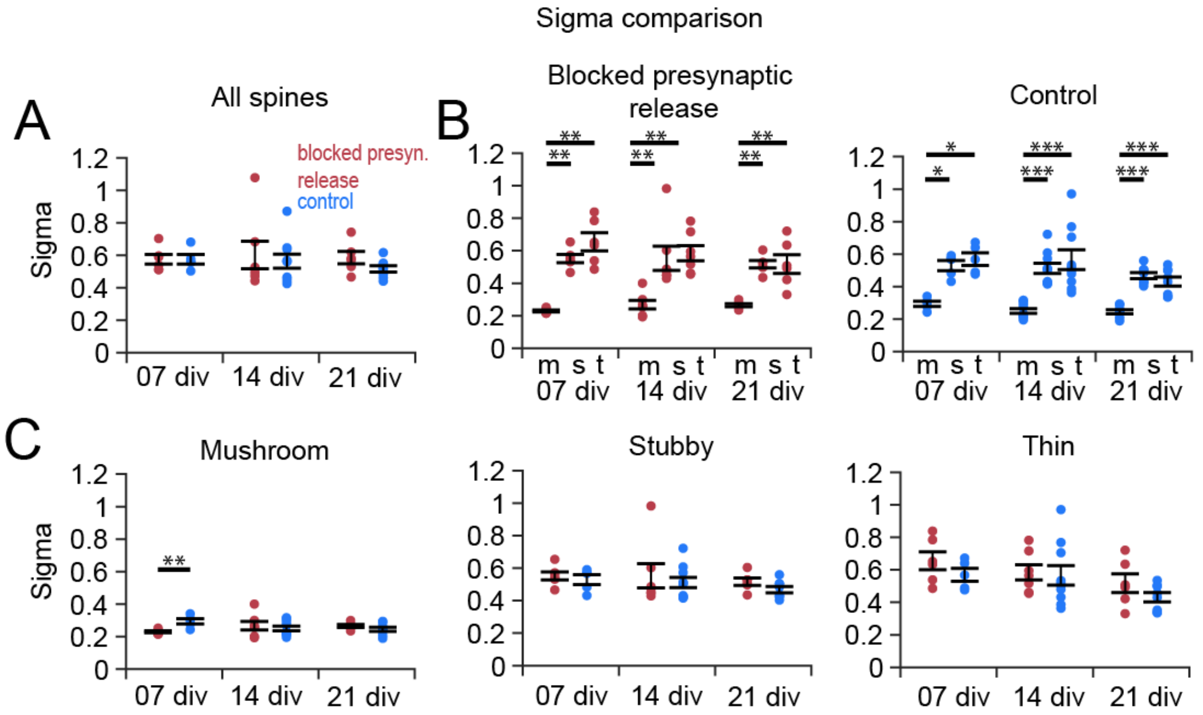


***Supplementary Figure 8. The sigma comparison in individual CA1 PCs from Munc-13 DKO (blocked presynaptic release) and WT (control) organotypic slice cultures, for all spine subtypes together and separately.*** *(A) The sigma values for all spine types were similar in the blocked presynaptic group and the control group. There are no significant differences between the groups or between the time points. The range of spine sizes stayed the same even when presynaptic release was blocked. (B) Comparing the sigma values of the different spine subgroups, in the blocked presynaptic release group (left panel) and the control group (right panel). Mushroom spines show the smallest values, meaning that they have the smallest range of spine sizes. (C) Analysis and comparison of sigma for each spine subtype separately. Mushroom spines (left panel) show the lowest overall sigma that stays unchanged through all cell ages and doesn't differ between the groups. Stubby (middle panel) and thin spines (right panel) show similar sigma values, however there are no significant differences between the two experimental groups in either spine type.*

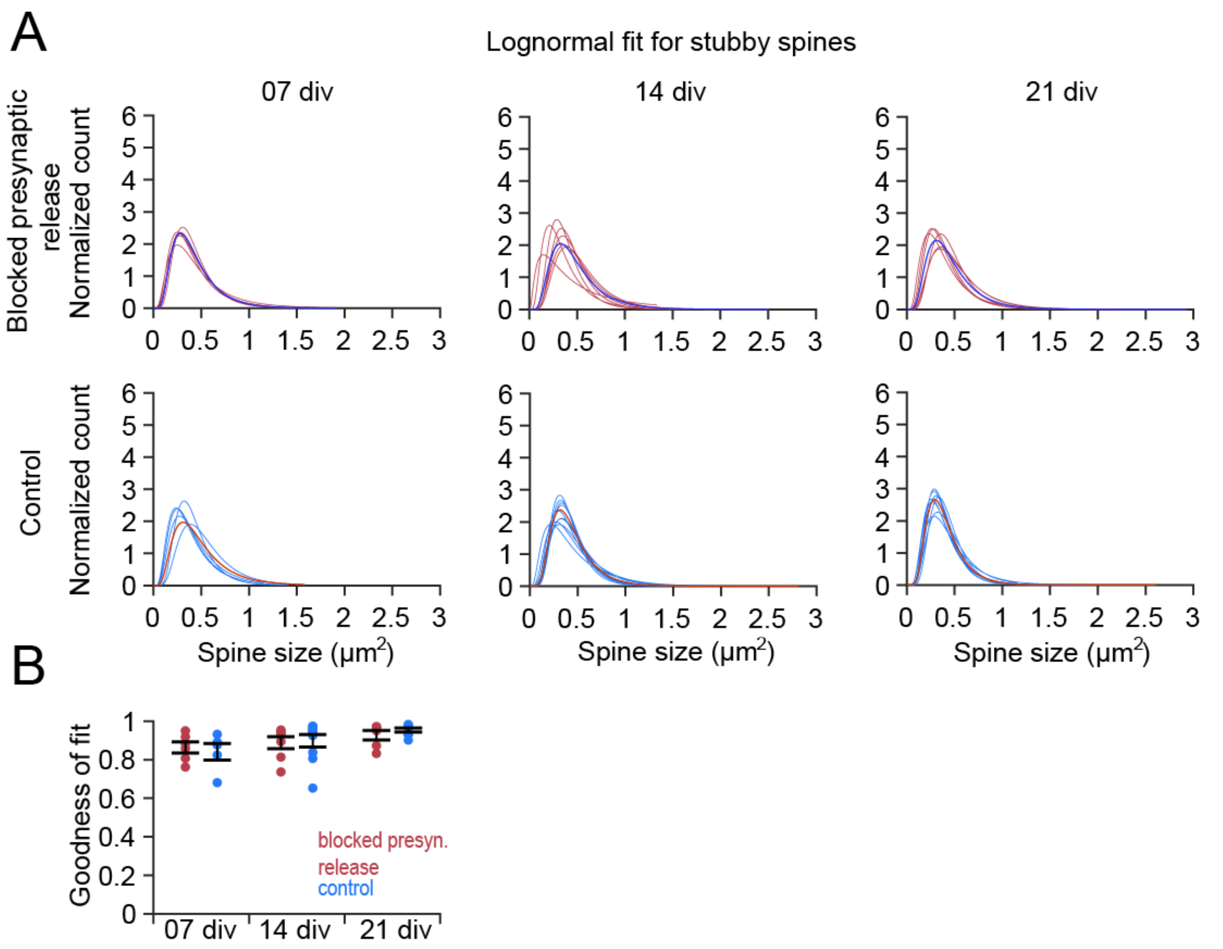


**Supplementary Figure 9.** ***Statistical analysis of stubby spines in individual CA1 PCs from Munc-13 DKO (blocked presynaptic release) and WT (control) organotypic slice cultures****. (A) Individual logfits of each cell in the blocked activity group and the control group for each time point separately. Each cell displays a lognormal-like distribution. (B) Analysis and comparison of goodness of fit values. No significant difference can be found between the two groups, but in the control group, the goodness of fit increases over time, from 07 div to 21 div ($p < 0.05$).*

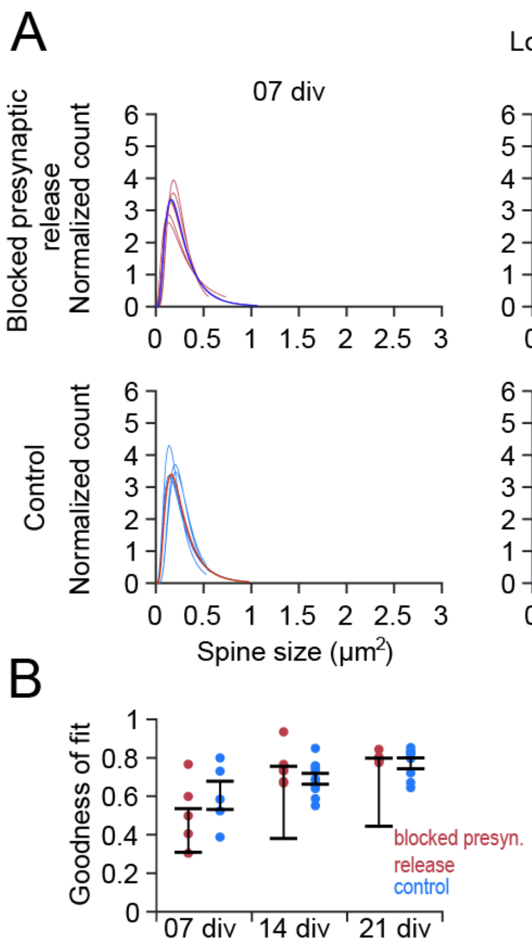


**Supplementary Figure 10.** *Statistical analysis of thin spines in individual CA1 PCs from Munc-13 DKO (blocked presynaptic release) and WT (control) organotypic slice cultures. (A) Individual logfits of each cell in the blocked activity group and the control group for each time point separately. Each cell displays a lognormal-like distribution. (B) Analysis and comparison of goodness of fit values. No significant differences can be found between the two groups, or in the time comparison.*

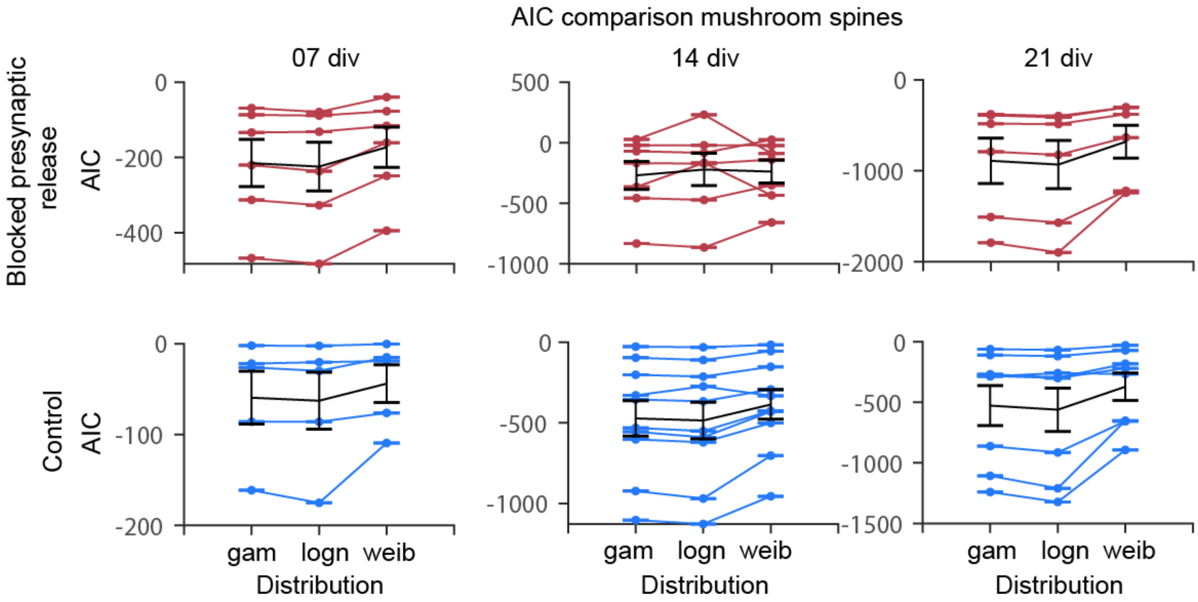


***Supplementary Figure 11. Akaike Information Criterion (AIC) in individual cells of the mushroom spine size data from CA1 PCs from Munc-13 DKO (blocked presynaptic release) and WT (control) organotypic slice cultures shows a small advantage of the lognormal distribution as the best model fit for the data compared to other skewed distributions.*** *At all time points, the lognormal distribution shows a slightly lower AIC value compare to other distributions. The lower the AIC, the better the fit of the model. In the cell specific analyses, in most comparisons, the lognormal distribution has the best fit. The only exception appears to be the mushroom spines at 14 div in the blocked presynaptic release group, where the lognormal distribution has slightly higher values. Each line represents one cell. gam – gamma distribution, logn – lognormal distribution, weib – Weibull distribution*
